# TRF2-mediated ORC recruitment underlies telomere stability upon DNA replication stress

**DOI:** 10.1101/2021.02.08.430303

**Authors:** Mitsunori Higa, Yukihiro Matsuda, Jumpei Yamada, Nozomi Sugimoto, Kazumasa Yoshida, Masatoshi Fujita

**Author notes:** Correspondence to: Kazumasa Yoshida, Department of Cellular Biochemistry, Graduate School of Pharmaceutical Sciences, Kyushu University, 3-1-1 Maidashi, Higashi-ku, Fukuoka 812-8582, Japan. Correspondence to: Masatoshi Fujita, Department of Cellular Biochemistry, Graduate School of Pharmaceutical Sciences, Kyushu University, 3-1-1 Maidashi, Higashi-ku, Fukuoka 812-8582, Japan.

## Abstract

Telomeres are intrinsically difficult-to-replicate regions of eukaryotic chromosomes. Telomeric repeat binding factor 2 (TRF2) binds to origin recognition complex (ORC) to facilitate the loading of ORC and the replicative helicase MCM complex onto DNA at telomeres. However, the biological significance of the TRF2-ORC interaction for telomere maintenance remains largely elusive. Here, we employed a separation-of-function TRF2 mutant with mutations in two acidic acid residues (E111A and E112A) that specifically inhibited the TRF2-ORC interaction in human cells without substantially inhibiting TRF2 interactions with its other binding partners. The TRF2 mutant was impaired in ORC recruitment to telomeres and showed increased replication stress-associated telomeric DNA damage and telomere instability. Furthermore, overexpression of an ORC1 fragment (amino acids 244–511), which competitively inhibited the TRF2-ORC interaction, increased telomeric DNA damage under replication stress conditions in human cells. Taken together, these findings suggest that TRF2-mediated ORC recruitment contributes to the suppression of telomere instability.

## Introduction

The origin recognition complex (ORC) is composed of 6 subunits (ORC1-6) and binds to replication origins distributed across the eukaryotic genome (Bleichert, 2019; Parker et al., 2017). Human ORC binds to origin DNA with no obvious sequence specificity and binding principally depends on the chromatin environment (Dellino et al., 2013; Higa et al., 2017a; Miotto et al., 2016; Parker et al., 2017; Picard et al., 2014). ORC-binding sites share several common characteristics, such as the presence of transcriptional start sites with an open chromatin structure, active histone modifications, and CpG islands (Dellino et al., 2013; Miotto et al., 2016; Picard et al., 2014). In addition, various chromatin-associated proteins, such as HP1α, dimethylated histone H4 (H4-K20me2), ORCA, and telomeric repeat binding factor 2 (TRF2) (Higa et al., 2017a; Parker et al., 2017), associate with the ORC complex and act as local ORC recruiters. In late M to G1 phase, ORC, and the additional licensing factors CDC6 and Cdt1, cooperatively promote the loading of minichromosome-maintenance (MCM) complex, a core component of the replicative helicase (Bleichert, 2019; Parker et al., 2017; Sugimoto and Fujita, 2017). During the following S phase, activated cyclin-dependent kinases (Cdks) and Dbf4-dependent kinase (DDK) trigger the initiation of DNA replication. Phosphorylation of MCM is a prerequisite for origin firing, while ORC, CDC6, and Cdt1 are downregulated by phosphorylation to prevent MCM re-loading and DNA re-replication (Fujita, 2006; Hills and Diffley, 2014). Replication stress-induced fork stalling activates MCMs pre-loaded onto dormant origins, promoting origin firing to assist in the completion of replication. Reduction in MCM levels causes DNA breaks, micronuclei formation, and genome instability, eventually leading to cellular senescence, inflammation and increased cancer risk (Chuang et al., 2010; Ibarra et al., 2008; Kawabata et al., 2011; Kunnev et al., 2010; McNairn et al., 2019; Sedlackova et al., 2020; Shima et al., 2007).

Telomeres are the terminal regions of linear chromosome. In mammals, the chromosome ends form telomere loops (T-loops), protecting DNA ends from detection by DNA damage response sensors (de Lange, 2018; Van Ly et al., 2018). End-protection is mostly achieved by telomere-specific chromatin-binding proteins that form the shelterin complex, comprised of TRF1, TRF2, RAP1, TIN2, TPP1, and POT1 (de Lange, 2018). DNA replication forks are prone to arrest and/or collapse at telomeres, leading to telomere instability, since telomeric higher-order structures and repetitive DNA sequences can interfere with fork progression (Gilson and Géli, 2007; Higa et al., 2017a; Martínez and Blasco, 2015; Pfeiffer and Lingner, 2013; Stroik and Hendrickson, 2020). In particular, guanine quadruplex (G4 DNA), DNA topological stress, and protective T-loop structures have been shown to lead to telomere instability if left unresolved during S phase (Fouché et al., 2006; Martínez et al., 2009; Sarek et al., 2015; Sfeir et al., 2009; Vannier et al., 2012). To facilitate telomere replication, the shelterin complex recruits additional factors to remove such obstacles during DNA replication. For example, TRF2 recruits Apollo, a nuclease that relieves topological stress (Chen et al., 2008; van Overbeek and de Lange, 2006; Ye et al., 2010); RTEL1 helicase, which dismantles the G4 DNA and the T-loop structure (Sarek et al., 2019, 2015; Vannier et al., 2012); and SLX4, a multitasking protein involved in the maintenance of telomere stability and the replication stress response (Guervilly and Gaillard, 2018; Wan et al., 2013). Overall, a complicated protein network is required to achieve efficient duplication of telomeric DNA tracts.

TRF2 is suggested to play a role in ORC and MCM loading at telomeres. TRF2 directly binds to ORC through the ORC1 subunit (Atanasiu et al., 2006; Higa et al., 2017b; Tatsumi et al., 2008) and RNA interference (RNAi)-mediated TRF2 silencing decreases loading of ORC and MCM onto telomeric DNA (Deng et al., 2007; Tatsumi et al., 2008), suggesting that replication origins are assembled at telomeres through the TRF2-ORC interaction. Indeed, DNA combing experiments have demonstrated replication initiation events occurring inside the telomeric tract (Drosopoulos et al., 2020, 2015, 2012). These initiation events may play an important role in telomere maintenance as the persistent arrest of replication forks within a telomere would otherwise result in under replication due to the absence of a converging fork (Alver et al., 2014). Considering the inherent difficulties associated with telomere replication, these telomeric replication origins may contribute to the complete duplication of telomeric tracts (Alver et al., 2014).

The biological role of the TRF2-ORC interaction is not fully understood, in part because siRNA-mediated depletion of TRF2 or essential ORC subunits inevitably affects other fundamental functions of these factors; for example, TRF2 knockdown affects telomere protection, while ORC1 knockdown compromises genome-wide DNA replication licensing. In this study, we evaluated the biological relevance of the TRF2-ORC interaction by two different means: firstly, by using a separation-of-function TRF2 mutant defective only in ORC binding, we show that the TRF2-ORC interaction promotes the recruitment of ORC and MCM at telomeres, and prevents telomere DNA damage and telomere instability under DNA replication stress conditions; secondly, we demonstrate that overexpression of an ORC1 fragment (amino acids 244-511), which binds to TRF2, competitively inhibits ORC recruitment at telomeres and induces the replication stress-associated telomere DNA damage in cells. These results strongly suggest that ORC recruitment by TRF2 underlies proficient telomere replication and telomere stability.

## Results

### TRF2 E111/E112 residues are required for efficient recruitment of ORC but not for interactions with other binding partners

To evaluate the TRF2-ORC interaction we sought to identify a separation-of-function TRF2 mutant defective only in its interaction with ORC. We previously reported that the dimerization of TRF2 is required for the recruitment of ORC (Higa et al., 2017b). Using the *lacO-LacI* protein tethering assay in U2OS 2-6-3 cells containing an array of 256 *lacO* repeats on chromosome 1 (Janicki et al., 2004), we have also shown that the TRFH (TRF homology) domain of TRF2 (amino acids 45–244) fused to LacI is sufficient for ORC and MCM recruitment to this *lacO* array (Higa et al., 2017b). TRF2 V52D and N53P mutations in the TRFH domain impairs TRF2 dimer formation and ORC recruitment by TRF2 (Higa et al., 2017b). However, the TRF2 (V52D/N53P) mutant is not suitable as a specific ORC interaction mutant because TRF2 dimerization is critical for its other functions, including telomere binding. To identify a specific TRF2-ORC interaction mutant, residues predicted to be required for binding to ORC1 were identified based on the crystal structure of the TRF2 TRFH domain (PDB code 4M7C) and the previously published data described above (Figure 1A). TRF2 residues were selected according to the following three criteria: (1) Since TRF2 dimerization is required for ORC1 binding and recruitment, we speculated that ORC1 recognizes a specific structure of the TRFH dimer created around the interface of the two monomers; hence, the residues selected were within helix 2 to helix 3 of the TRFH located in proximity to the dimer interface; (2) selected residues were exposed on the surface of the protein; (3) selected residues were not conserved in TRF1, since TRF1 is unable to recruit ORC (Higa et al., 2017b). Fourteen residues (Y73, G74, V88, P90, K93, E94, H95, T96, S98, R102, E111, E112, S119, and M122) met these criteria (Figure 1A), and six TRF2 mutants were generated by substituting the selected residues with alanine (Y73A/G74A, V88A/P90A, K93A/E94A/H95A/T96A, S98A/R102A, E111A/E112A, and S119A/M122A; referred to as YG, VP, KEHT, SR, EE and SM respectively).

**Figure 1.**
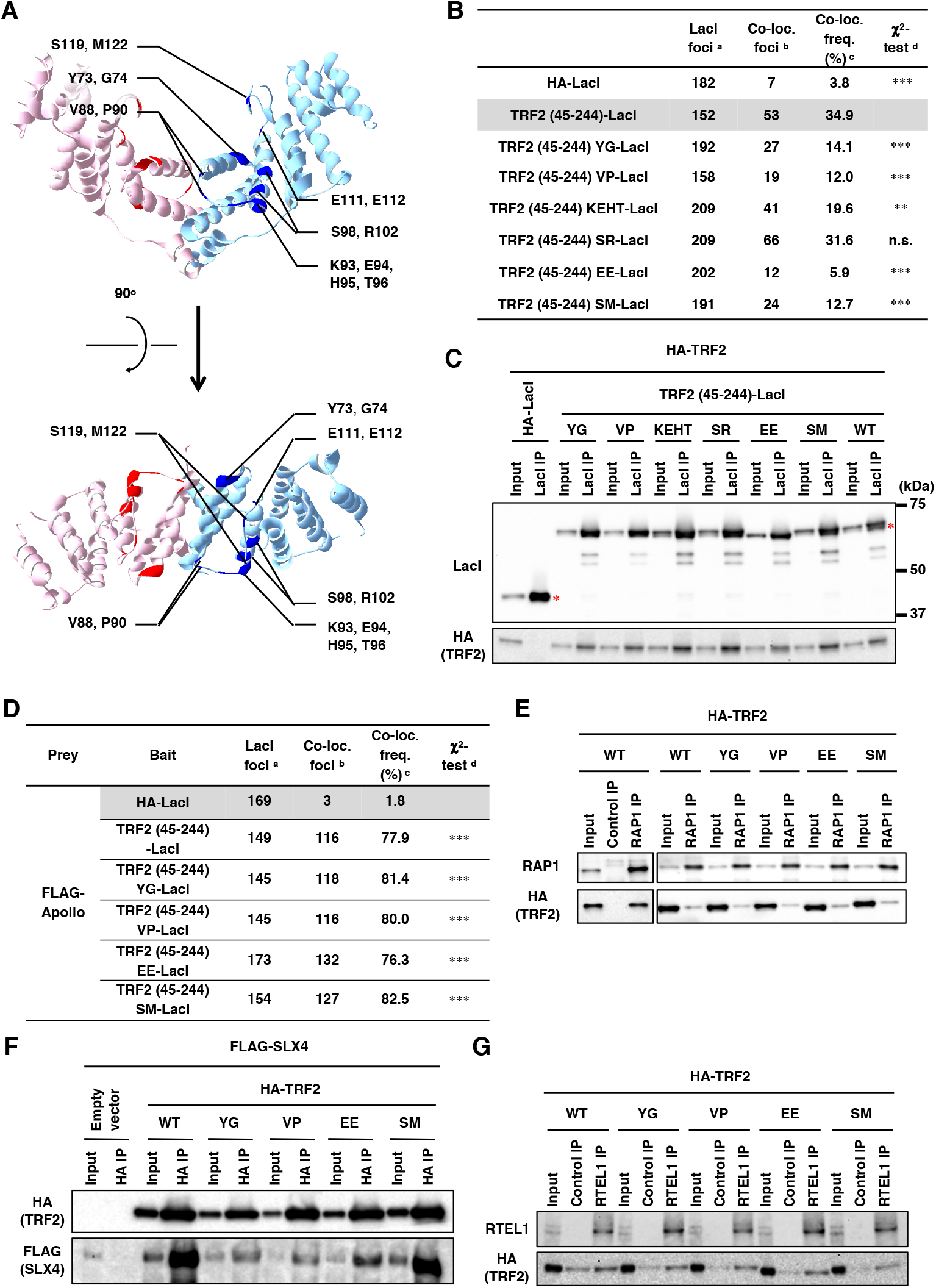
Generation and characterization of a series of TRF2 mutants with alanine-substitutions in the TRFH domain. **(A)** Crystal structure of the TRF2 TRFH domain (PDB code 4M7C). Amino acid residues mutated in this study are highlighted and individual TRF2 monomers are depicted with cyan and pink ribbons. The 14 selected residues are marked with blue or red. *Top,* front view. *Bottom,* top view. **(B)** U2OS 2-6-3 cells were transfected with HA-LacI, TRF2 (45–255)-LacI, or alanine-substitution mutant TRF2 (45–244)-LacI (YG, VP, KEHT, SR, EE, and SM) for 24 h, and then co-immunostained with anti-LacI antibody and anti-ORC1 antibody, followed by DAPI counterstaining. Co-localization frequencies of ORC1 with LacI proteins were examined. The values represent the sum scores from two biologically independent experiments. ^a^ Total number of LacI foci. ^b^ Number of LacI foci co-localizing with ORC1 foci. ^c^ Co-localization frequency of ORC1 foci with LacI foci. ^d^ Results of the χ^2^ test vs. TRF2 (45–244)-LacI. **, P < 0.01; ***, P < 0.001; n.s., not significant. **(C)** U2OS cells were co-transfected with the indicated expression vectors. At 42 h post-transfection, cells were harvested and subjected to immunoprecipitation with anti-LacI antibody. Immunoprecipitates (IPs) and 3% of the input were separated by SDS-PAGE, followed by immunoblotting with the indicated antibodies. Asterisks indicate the position of full-length LacI-fused proteins. WT, wild-type. **(D)** U2OS 2-6-3 cells were co-transfected with the indicated expression vectors for 24 h. Co-localization frequency of FLAG-Apollo and LacI-fused proteins was examined. The values represent the sum scores from at least two biologically independent experiments. ^a^ Total number of LacI foci. ^b^ Number of LacI foci colocalizing with FLAG-Apollo foci. ^c^ Co-localization frequency of FLAG-Apollo foci with LacI foci. ^d^ Results of the χ^2^ test vs. HA-LacI. ***, P < 0.001. **(E)** At 42 h posttransfection with the indicated vectors, HCT116 cells were harvested and immunoprecipitated with an anti-RAP1 antibody or normal rabbit IgG (Control IP). IPs and 3% of input were analyzed by immunoblotting with the indicated antibodies. **(F)** At 42 h post-transfection with the indicated vectors, HEK293T cells were harvested and immunoprecipitated with an anti-HA antibody. IPs and 0.5% of input were analyzed by immunoblotting with the indicated antibodies. **(G)** HEK293T cells were cotransfected with the indicated expression vectors for 42 h, cross-linked with formaldehyde and then solubilized. Soluble fractions were immunoprecipitated with an anti-RTEL1 antibody or normal mouse IgG (Control IP). IPs and 2% of input for RTEL1 or 0.1% of input for HA were analyzed by immunoblotting with the indicated antibodies.

The ability of the mutants to recruit ORC was examined using the *lacO-LacI* assay (Figure 1B). The results clearly showed that TRF2 (45-244) EE-LacI was defective in recruiting ORC1 to the *lacO* array. In addition, YG, VP, KEHT, and SM mutants showed partially decreased co-localization with ORC1, suggesting that these residues are also involved in the interaction with ORC. Immunoprecipitation assays demonstrated that all the prepared mutants of TRF2 (45-244)-LacI can bind to HA-TRF2 with similar affinity to that of wild-type TRF2 (45-244)-LacI (Figure 1C), indicating that these mutations do not affect dimerization. TRF2 functions as a hub protein that directly binds to telomeres and recruits various factors required for telomere maintenance. For example, a component of shelterin complex, RAP1, binds to telomeres concomitant with TRF2 (Li et al., 2000; Sfeir et al., 2010). In addition, several factors such as Apollo, SLX4, and RTEL1 access telomeres by binding to the TRF2 TRFH domain, thereby facilitating telomeric DNA replication (Higa et al., 2017a; Martínez and Blasco, 2015). Therefore, we next examined whether our TRF2 mutants were proficient in binding to these partners (Figure 1D-G). YG, VP, EE, and SM mutants recruited Flag-Apollo to the *lacO* array to a similar extent as wild-type TRF2 (Figure 1D). These mutants could also bind RAP1 (Figure 1E), SLX4 (Figure 1F), and RTEL1 (Figure 1G) in immunoprecipitation assays. Collectively, these results suggest that TRF2 EE is a specific TRF2 mutant defective in ORC recruitment alone.

### Recruitment of ORC and MCM to telomeres is inhibited by the TRF2 E111A/E112A mutation in HeLa cells

To evaluate the effects of substitution of endogenous wild-type TRF2 with the TRF2 EE mutant, we established HeLa clones carrying EE mutations in the *TERF2* gene by gene-editing using CRISPR-Cas9 and single stranded oligonucleotide DNA (ssODN) encoding the EE mutations and a diagnostic NruI restriction site (Figure 2A-D). Gene-editing with an ssODN that introduced the NruI site alone was performed in parallel to establish a control clone (Figure 2A, ssODN WT). Restriction fragment length polymorphism (RFLP) analysis showed that TRF2 WT 7-1, TRF2 EE 4-3, and TRF2 EE 7-7 clones have at least one *TERF2* allele carrying the mutated sequences with the newly introduced NruI site (Figure 2B). The target sites in the *TERF2* gene exon 2 were amplified and sequenced, as described in Materials and Methods, and no wild-type sequences or in-frame insertions/deletions were detected (Figure 2C). In all TRF2-edited clones, TRF2 protein levels were decreased to 20–40% of that in the parental HeLa cell line (Figure 2D). It is possible that this is caused by frameshift mutations in other TRF2 alleles. To examine whether TRF2 EE affects the recruitment of ORC to telomeres, ChIP-qPCR assays to examine telomere-specific chromatin loading were performed using an anti-FLAG antibody in TRF2-edited clones expressing ORC1-3×FLAG (Figure 2E and Figure 2– figure supplement 1), and telomere DNA enrichment relative to the *LMNB2* replication origin (Giacca et al., 1994; Sugimoto et al., 2011) was calculated (Figure 2E). ORC1 binding to telomeres was decreased to approximately 60% in the WT clone compared with the parental HeLa cells. This decrease may be associated with the reduced TRF2 protein levels in the WT clone. In the two EE clones, ORC1 binding to telomeres was decreased 4-fold compared with the WT clone. This impairment was not attributable to the expression level of ORC1 −3×FLAG (Figure 2F) or to cell cycle distribution (Figure 2G). We next examined whether MCM recruitment to telomeres was affected in the TRF2-edited clones using MCM7 ChIP-qPCR. MCM7 binding to telomeres was significantly decreased in the two EE clones when compared with the WT clone (Figure 2H), while MCM7 DNA binding in general was unaffected (Figure 2I). These data suggest that TRF2-ORC binding is critical for the recruitment of ORC and MCM to telomeres.

**Figure 2.**
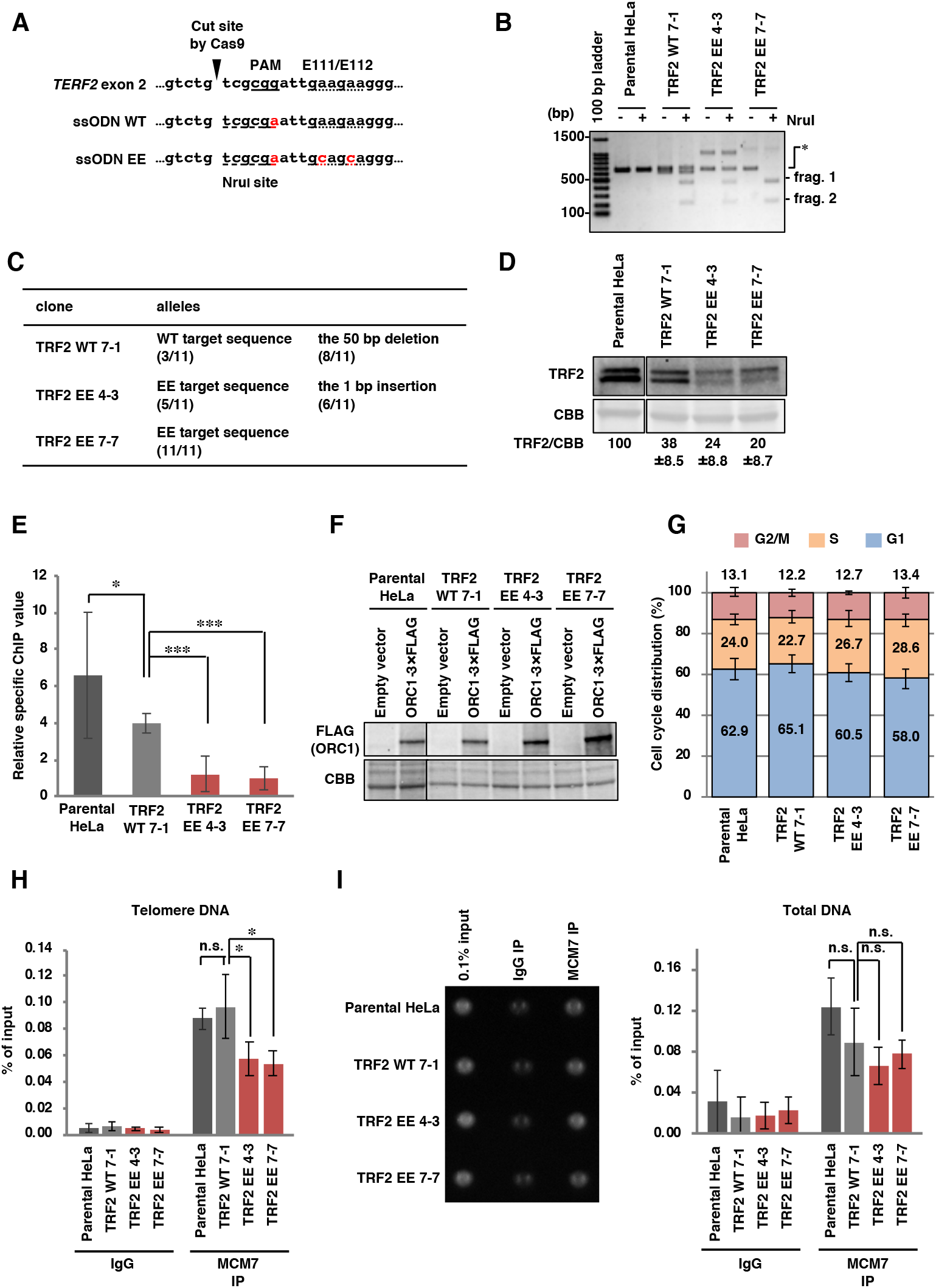
ORC recruitment to telomeres is impaired in TRF2 EE clones. **(A)** The sequences of single stranded oligo DNA (ssODN) used to establish TRF2-edited HeLa clones. *Top,* reference sequence of the target site. *Middle,* ssODN used for the TRF2 WT clone. *Bottom,* ssODN used for TRF2 EE clones. Introduced mutations are shown in red. The PAM sequence required for Cas9 recognition is underlined. Codons corresponding to the E111/E112 residues are marked with a dotted underline. The newly introduced NruI site is marked with a dashed underline. **(B)** Restriction fragment length polymorphism analysis of the HeLa clones. The target site was amplified from genomic DNA derived from each cell line by PCR. Amplicons were incubated with or without NruI restriction enzyme and then separated by agarose gel electrophoresis, followed by EtBr staining. When the target genes were edited, NruI treatment produced 427 bp and 209 bp fragments (frag. 1 and frag. 2, respectively) from the 636 bp PCR product (*). **(C)** Summary of the sequencing of the target sites in the *TERF2* gene of each cell line. For detail, see Materials and Methods. Eleven colonies from each clone were analyzed. **(D)** Whole cell extracts were separated by SDS-PAGE, and then analyzed by immunoblotting with anti-TRF2 antibody or by CBB staining (as a loading control). The signal intensities of TRF2 and CBB bands were quantified, and the TRF2/CBB signal ratio is shown relative to that of parental HeLa cells. The means ± SDs are shown (n = 3). **(E-F)** At 48 h post-transfection with empty vector or ORC1-3 × FLAG expression vector, cells were subjected to ChIP with an anti-FLAG antibody. (E) Purified DNA from input and immunoprecipitates was analyzed by qPCR with primer pairs amplifying either a telomeric sequence or the *LMNB2* replication origin. Relative specific ChIP values were calculated as follows: (Telomeric DNA as a % of input from ORC1-3 × FLAG ChIP-Telomeric DNA as a % of input from empty vector ChIP) / *(LMNB2* origin DNA as a % of input from ORC1-3 × Flag ChIP - *LMNB2* origin DNA as a % of input from empty vector ChIP). The means ± SDs are shown (n = 9). * P < 0.05; ***, P < 0.001 (two-tailed Student’s *t*-test). (F) Lysates were separated by SDS-PAGE and then analyzed by immunoblotting with anti-FLAG antibody and by CBB staining (as a loading control). **(G)** Cells were harvested and stained with propidium iodide to allow cell cycle analysis by flow cytometry. The means ± SDs are shown (n = 3). **(H and I)** Cells were subjected to ChIP assay with control IgG or anti-MCM7 antibody. (H) Purified DNAs from input and immunoprecipitates were analyzed by qPCR with a primer pair amplifying a telomeric sequence. Results are shown as the percent of input DNA. The means ± SDs are shown (n = 4). *, P < 0.05; n.s., not significant (two-tailed Student’s t-test). (I) *Left,* the total amounts of co-precipitated DNA and input DNA were analyzed using SYBR Gold staining and UV photography. *Right,* results are shown as the percent of input DNA. The means ± SDs are shown (n = 4). n.s., not significant (two-tailed Student’s t-test).

**Figure 2–figure supplement 1.**
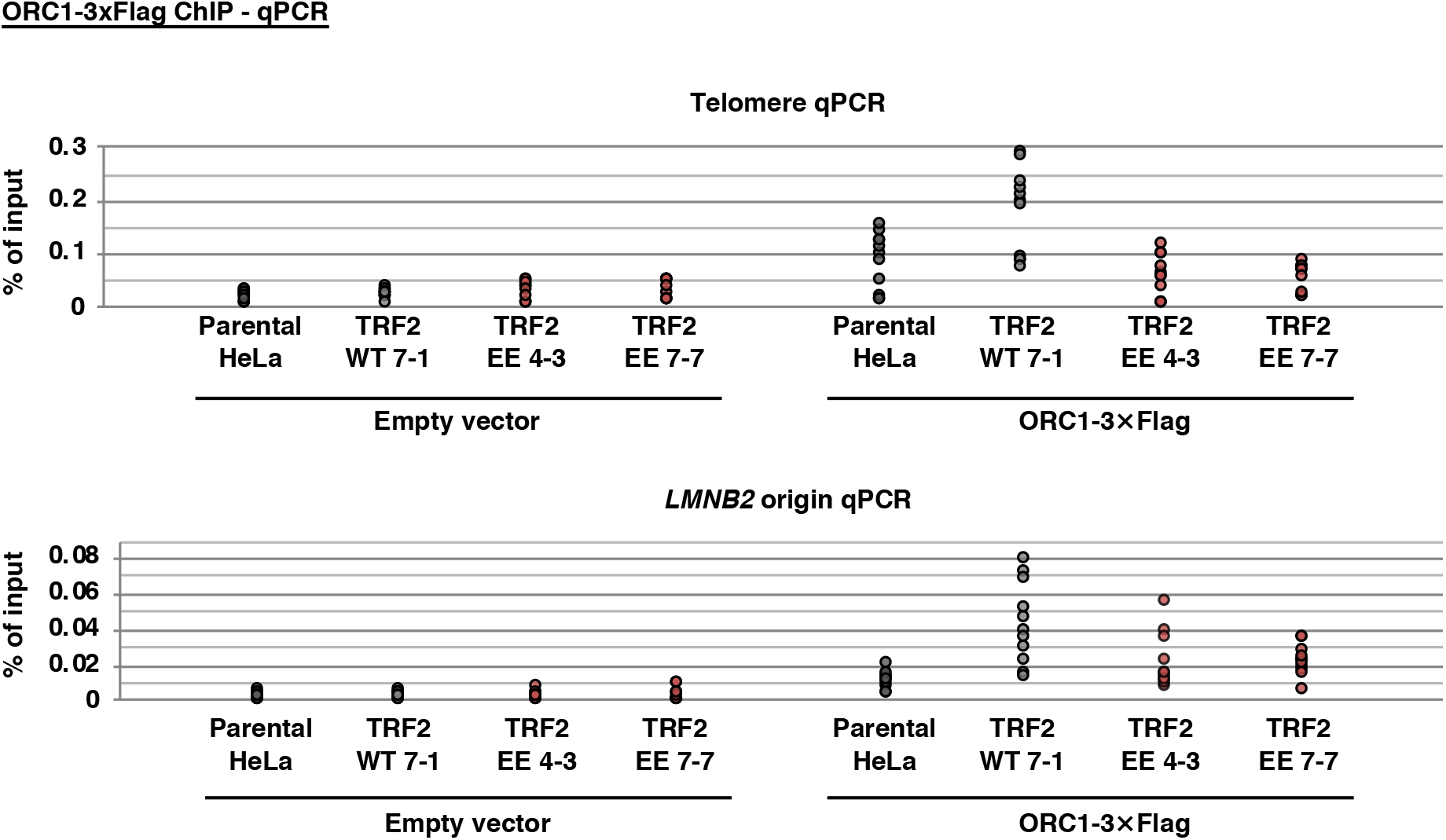
Source data of ChIP experiments. % of input values used for the calculation in Figure 2E are shown.

### Telomere maintenance is impaired in HeLa TRF2 E111A/E112A clones under DNA replication stress conditions

The phenotype of TRF2-edited clones was analyzed with particular focus on telomere maintenance under DNA replication stress conditions. First, we investigated telomere DNA damage. After exposing cells to 0.1 mM hydroxyurea (HU), telomeres and 53BP1, a marker for DNA double-strand breaks (Panier and Boulton, 2014), were visualized by fluorescence in situ hybridization (FISH) with a peptide nucleic acid (PNA) probe targeting a telomere-specific sequence, and immunofluorescence staining, respectively (Figure 3A and B). In the absence of HU-induced replication stress, the number of telomere dysfunction-induced foci (TIFs), detected as telomeric foci co-localizing with 53BP1 (d’Adda di Fagagna et al., 2003; Takai et al., 2003), increased in the WT clone (Figure 3B, Parental HeLa/DDW vs. WT 7-1/DDW). This increase is possibly related to the decreased TRF2 protein level in WT 7-1 (see Figure 2D), which may lead to insufficient telomere protection (Celli and de Lange, 2005; van Steensel et al., 1998). In the absence of HU, the number of endogenous TIFs in the two EE clones were comparable to the WT clone, suggesting that there was no increase in the basal level of telomere damage in the EE clones. Importantly, HU treatment increased the number of TIFs in the two EE clones but not in the WT clone, indicating a defect in the response to HU-induced telomere replication stress in the EE clones.

**Figure 3.**
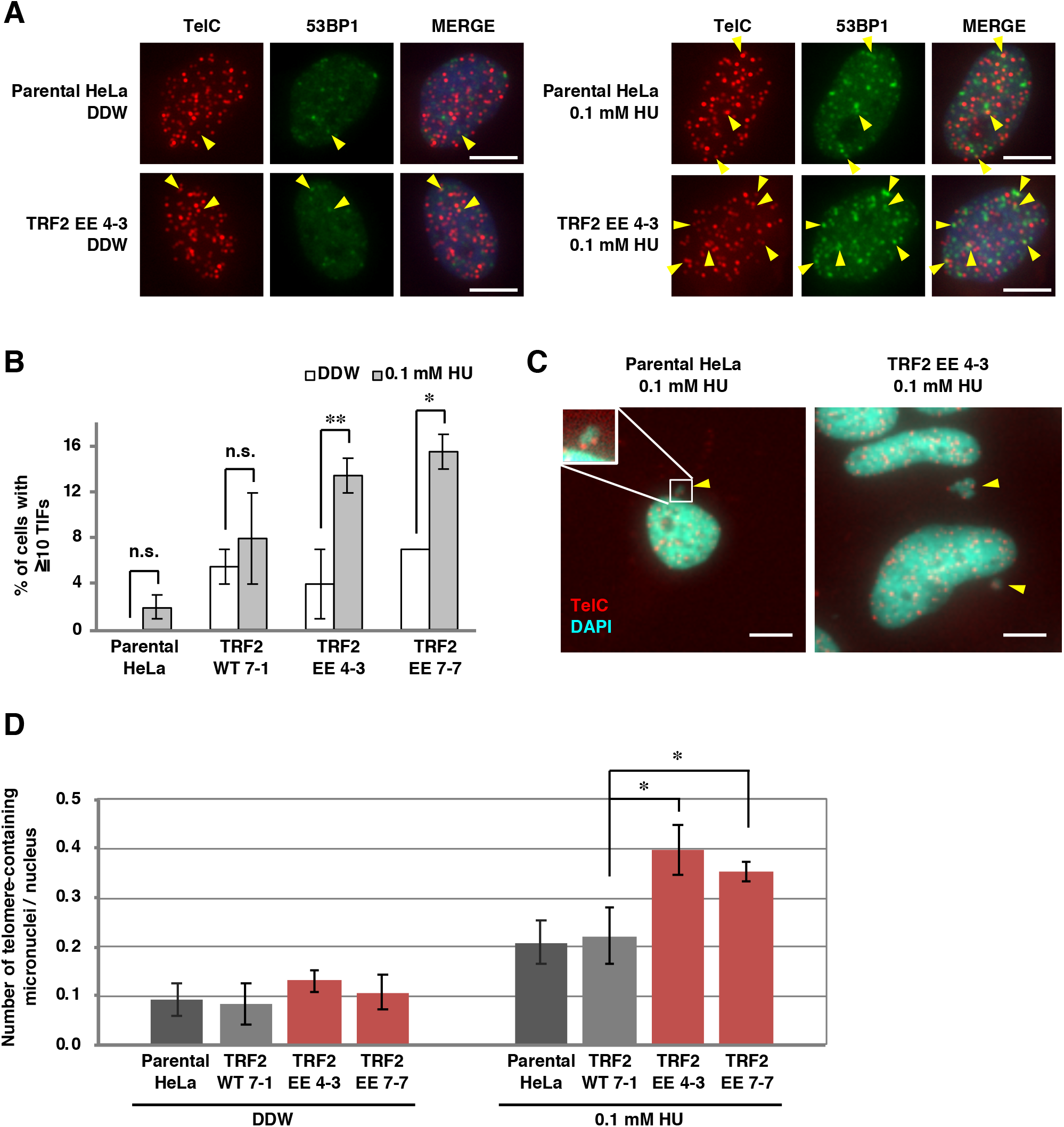
Telomere instability in TRF2 EE clones upon DNA replication stress. **(A and B)** Cells treated with 0.1 mM hydroxyurea (HU) for 16 h were stained with PNA FISH using a telomere probe (TelC, red), followed by immunostaining with anti-53BP1 antibody (green) and DAPI counterstaining (blue). (A) Representative images; telomere foci co-localizing with 53BP1 (53BP1 TIFs) are indicated by yellow arrowheads. Scale bar, 10 mm. (B) The frequencies of cells carrying ≥10 53BP1 TIFs were quantified. The means ± SDs are shown (n = 2). One hundred cells were analyzed for each sample. The sum scores are used for statistical analyses. n.s., not significant; *, P < 0.05; **, P < 0.01 (Fisher’s exact test). **(C and D)** Cells treated with 0.1 mM HU for 72 h were stained with PNA-FISH using a telomere probe (red), followed by DAPI counterstaining (cyan). (C) Representative images; telomere-containing micronuclei are indicated by yellow arrowheads. Scale bar, 10 mm. (D) The number of nuclei and telomere-containing micronuclei were counted, and the ratio of telomere-containing micronuclei/nucleus was calculated. The means ± SDs are shown (n = 3). At least 82 cells were analyzed for each sample. *, P < 0.05 (two-tailed Student’s *t*-test).

Defective DNA replication leads to micronuclei formation (Chan et al., 2009; Ying et al., 2013); therefore, we next measured the number of micronuclei containing telomeric FISH signals (telomere-containing micronuclei) as an index of telomere instability (Lindberg et al., 2008) (Figure 3C and D). In the absence of HU, the frequency of telomere-containing micronuclei did not differ between the cell lines tested. However, HU induced significantly more telomere-containing micronuclei in EE clones than the WT clone (approximately 2-fold). Taken together, these results suggest that TRF2-mediated recruitment of ORC to telomeres contributes to the maintenance of telomere stability under DNA replication stress conditions.

### A broad region of ORC1 including amino acids 411–510 is required for the recruitment by TRF2

As an alternative approach to evaluate the physiological significance of the TRF2-ORC interaction, we searched for a specific ORC1 mutant specifically defective for TRF2 binding alone. Several proteins that bind to the TRFH domain of TRF2, such as Apollo and SLX4, share a TRF2 binding motif (F/Y/HxLxP) (Chen et al., 2008; Wan et al., 2013). The leucine residues in the binding motifs of Apollo and SLX4 are important for their interaction with TRF2 (Chen et al., 2008; Wan et al., 2013). This short motif is similarly found in human ORC1 at amino acid 227–231 (HTLTP). However, an ORC1 L229A mutant could be recruited by TRF2 in the *lacO-LacI* assay (Figure 4–figure supplement 1), suggesting that, unlike Apollo and SLX4, this motif in ORC1 may not be involved in TRF2 binding.

Next, we generated a series of ORC1 truncation mutants (Figure 4A) and examined their recruitment by TRF2 (45–244)-LacI (Figure 4B). Transiently-expressed FLAG-ORC1 (2–511) frequently co-localized with TRF2 (45–244)-LacI (73%), indicating that the C-terminal AAA+ (ATPases associated with diverse cellular activities) and WH (winged-helix) domains are dispensable for binding to TRF2. By contrast, three further ORC1 truncation mutants (2–325, 2–244, and 2–85) were not recruited by TRF2 (45–244)-LacI, suggesting that a region spanning amino acid 326–511 of ORC1 is required for TRF2 binding. In addition, ORC1 (244–511), but not ORC1 (325-511), co-localized more frequently with TRF2 (45–244)-LacI (~90% and ~46%, respectively) than ORC1 (2– 511) (73%). These data suggest that ORC1 efficiently binds to TRF2 via a relatively broad region that includes amino acids 244–511. This is consistent with a previous report that GST-ORC1 (201– 511) interacts with TRF2 in HeLa cell extracts (Atanasiu et al., 2006).

**Figure 4.**
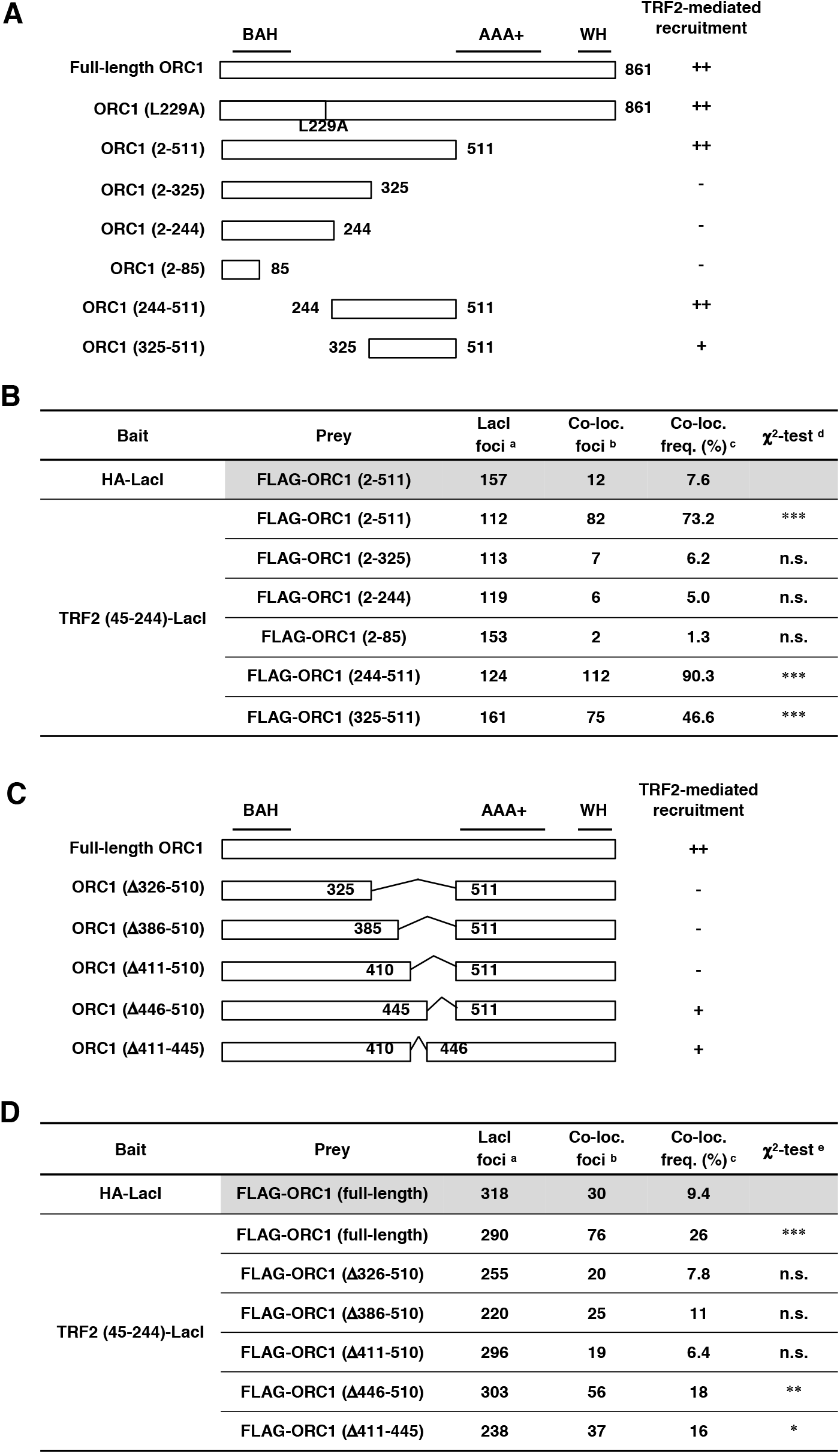
ORC1 amino acids 411-510 are required for efficient ORC1 recruitment by TRF2. **(A and C)** Schematics of ORC1 mutants used in this study. Domain information is shown at the top. A summary of their ability to bind TRF2 is shown on the right (see Figure 4B, D and Expanded View Figure EV2 for detail). BAH, bromo adjacent homology domain; AAA+, ATPase associated with diverse cellular activities domain; WH, winged helix domain. **(B and D)** Summary of the co-localization frequencies of ORC1 mutants with TRF2 (45-244)-LacI. U2OS 2-6-3 cells were co-transfected with the indicated ORC1 and LacI expression vectors. Co-localization frequency was examined as in Figure 1B. The values represent the sum score from at least two biologically independent experiments. ^a^ Total number of LacI foci. ^b^ Number of LacI foci co-localizing with FLAG foci. ^c^ Co-localization frequency of FLAG foci with LacI foci. ^d^ Results of the χ^2^ test vs. co-localization of HA-LacI and FLAG-ORC1 (2-511). ^e^ Results of the χ^2^ test vs. co-localization of HA-LacI and FLAG-ORC1 (full-length). *, P < 0.05; **, P < 0.01; ***, P < 0.001; n.s., not significant.

To further narrow the region required for TRF2 binding, we analyzed the recruitment of full-length ORC1 mutants lacking amino acids within the 326–510 region (Figure 4C and D). TRF2 (45– 244)-LacI recruited N-terminal FLAG-tagged ORC1 (full-length) at a significantly higher frequency (26%) than the control HA-LacI (9%). However, ORC1 (Δ326–510), ORC1 (Δ386–510), and ORC1 (Δ411–510) co-localized with TRF2 (45–244)-LacI at low frequencies, indicating that amino acids 411-510 of ORC1 are necessary for the recruitment by TRF2. By contrast, ORC1 (Δ446-510) and ORC1 (Δ411-445) partially retained its ability to bind to TRF2. These results suggest that multiple residues distributed across the 411-510 region are responsible for binding to TRF2. Because the loss of such a broad region is likely to have adverse effects on ORC1 stability and function, we considered ORC1 (Δ411-510) to be an unsuitable candidate for a specific ORC1 mutant defective only in TRF2 binding.

**Figure 4–figure supplement 1.**
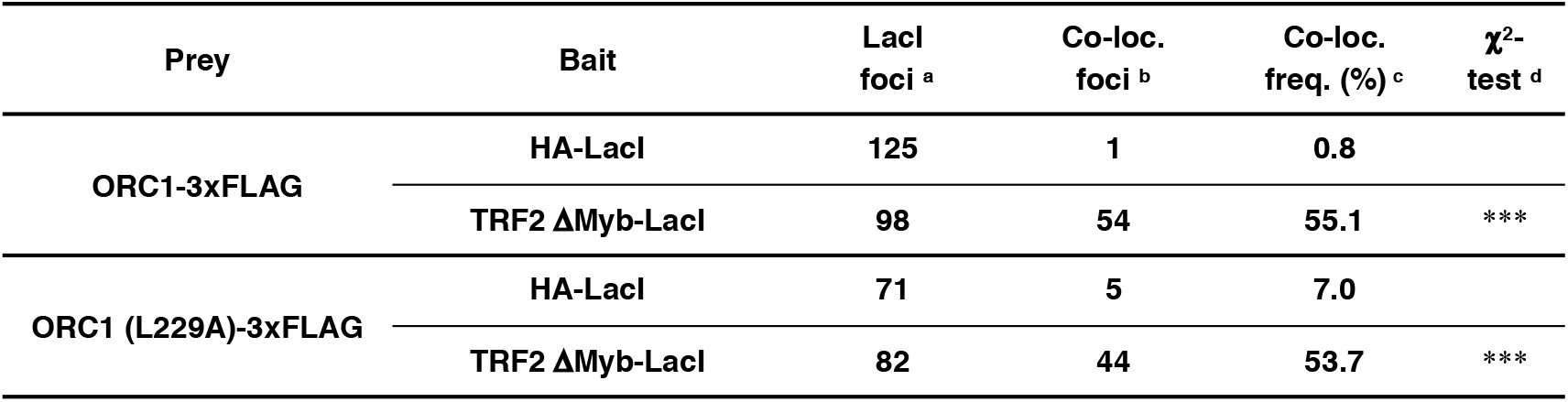
Co-localization frequencies of wild-type and L229A ORC1 with TRF2△Myb-LacI. After co-transfection with the indicated ORC1 and LacI expression vectors, U2OS 2-6-3 cells were subjected to co-immunostaining with anti-LacI and anti-FLAG antibodies. Co-localization frequency was examined. The values represent the score from a single experiment. ^a^ Total number of LacI foci. ^b^ Number of LacI foci colocalizing with FLAG foci. ^c^ Co-localization frequency of FLAG foci with LacI foci. ^d^ Results of the χ^2^ test in comparison with HA-LacI. ***, P < 0.001. TRF2ΔMyb, consisting of the amino acids 1-445 but lacking the Myb domain, has been reported to recruit ORC1 to the *lacO* array when fused to LacI (Higa et al. 2017 BBA Mol Cell Res 1864: 191-201). The data presented show that the co-localization frequency of ORC1 (L229A)-3 × FLAG with TRF2DMyb-LacI was ~54%, which was comparable to that of wild-type ORC1-3 × FLAG (~55%).

### Overexpression of an ORC1 (244–511) fragment competitively inhibits TRF2-ORC binding but does not affect the general function of ORC in the initiation of DNA replication

Since the TRF2 binding sites reside within amino acids 244 and 511 of ORC1, we examined whether overexpression of an ORC1 (244-511) construct could competitively inhibit the TRF2-ORC interaction (Figure 5). As shown in Figure 5A, FLAG-ORC1 (244-511) bound to TRF2 and competitively inhibited co-immunoprecipitation of full-length ORC1 with TRF2. In addition, overexpression of HA-ORC1 (244-511) impaired the TRF2 (45-244)-LacI-mediated recruitment of ORC1 to the *lacO* array (Figure 5B). Next, we examined whether ORC1 (244-511) overexpression affects the general ORC function during global DNA replication. Although MCM loading is tightly regulated to prevent DNA re-replication, simultaneous deregulation of licensing factors (Cdt1+ORC1 or Cdt1+CDC6) can induce re-replication in HEK293T cells, which is detected by FACS as DNA content higher than 4N (Sugimoto et al., 2009). We used this assay to examine the genome-wide functions of ORC in the presence of FLAG-ORC1 (244-511) (Figure 5C-E). Consistent with a previous report (Sugimoto et al., 2009), co-expression of Cdt1+ORC1 induced significant re-replication (~two-fold higher than that induced by expression of Cdt1 alone) (Figure 5C-E). This re-replication was dependent on ORC1 ATPase activity, as a mutation in the Walker B motif (D620A) abolished the induction of re-replication (Figure 5-figure supplement 1). Importantly, re-replication was induced by Cdt1 + ORC1 to a similar extent in the presence and absence of FLAG-ORC1 (244-511) (Figure 5C-E). These results suggest that overexpression of ORC1 (244-511) may specifically inhibit TRF2-ORC interaction with little effect on the genome-wide replication activity of ORC.

**Figure 5.**
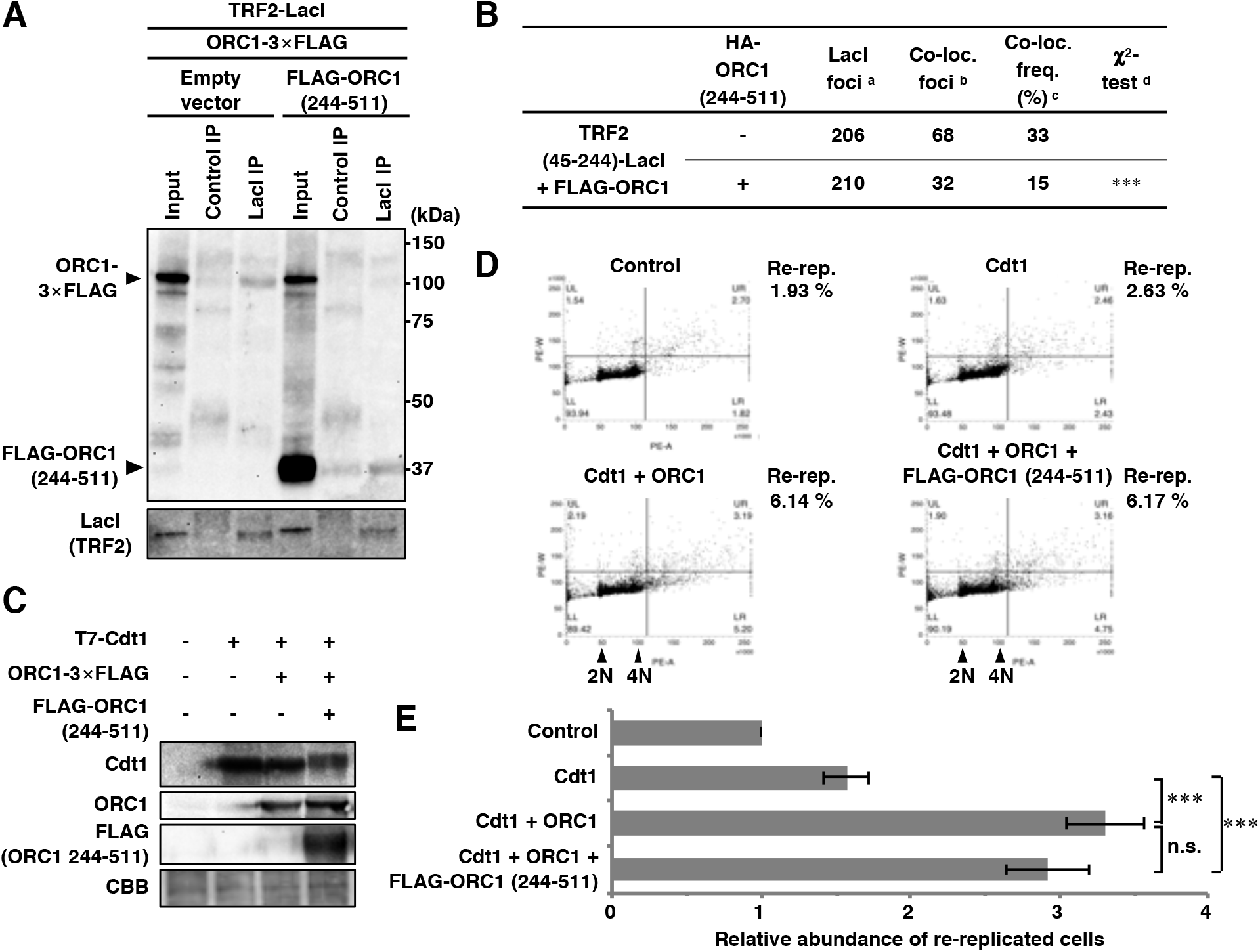
Overexpression of ORC1 (244-511) fragment competitively inhibits TRF2-ORC1 binding, but has little effect on the genomewide replication activity of ORC1. **(A)** HEK293T cells were co-transfected with the indicated expression vectors (equal amounts of each plasmid) for 42 h. After cross-linking with formaldehyde and solubilization, cell lysates were immunoprecipitated with anti-LacI antibody or normal mouse IgG (Control). IPs and 1% of the input were immunoblotted with the anti-FLAG or anti-LacI antibody. The data are representatives of two independent experiments. **(B)** U2OS 2-6-3 cells were co-transfected with the indicated expression vectors (equal amounts of each plasmid) for 24 h, then double-immunostained with the indicated antibodies, followed by DAPI staining. Co-localization frequencies of FLAG-ORC1 foci with LacI foci were examined. The values represent the sum score of two biologically independent experiments. ^a^ Total number of LacI foci. ^b^ Number of LacI foci co-localizing with FLAG-ORC1 foci. ^c^ Co-localization frequency of FLAG-ORC1 foci with LacI foci. ^d^ Results of the χ^2^ test vs. co-localization of FLAG-ORC1 and TRF2 (45-244)-LacI in the absence of HA-ORC1 (244-511). ***, P < 0.001. **(C-E)** HEK293T cells were co-transfected with mixture of the expression vectors (equal amounts of each plasmid) or their empty vectors (-) as indicated for 48 h. (C) Whole-cell lysates were subjected to immunoblotting with the indicated antibodies. CBB serves as a loading control. (D) DNA content was analyzed by flow cytometry. After excluding the sub-G1 fraction, the percentages of re-replicated cells (DNA content greater than 4N) were calculated (shown on the right). (E) The relative abundance of re-replicated cells as a proportion of control cells. The means and SDs from three biologically independent experiments were shown. ***, P < 0.001; n.s., not significant (Tukey-Kramer test).

**Figure 5–figure supplement 1.**
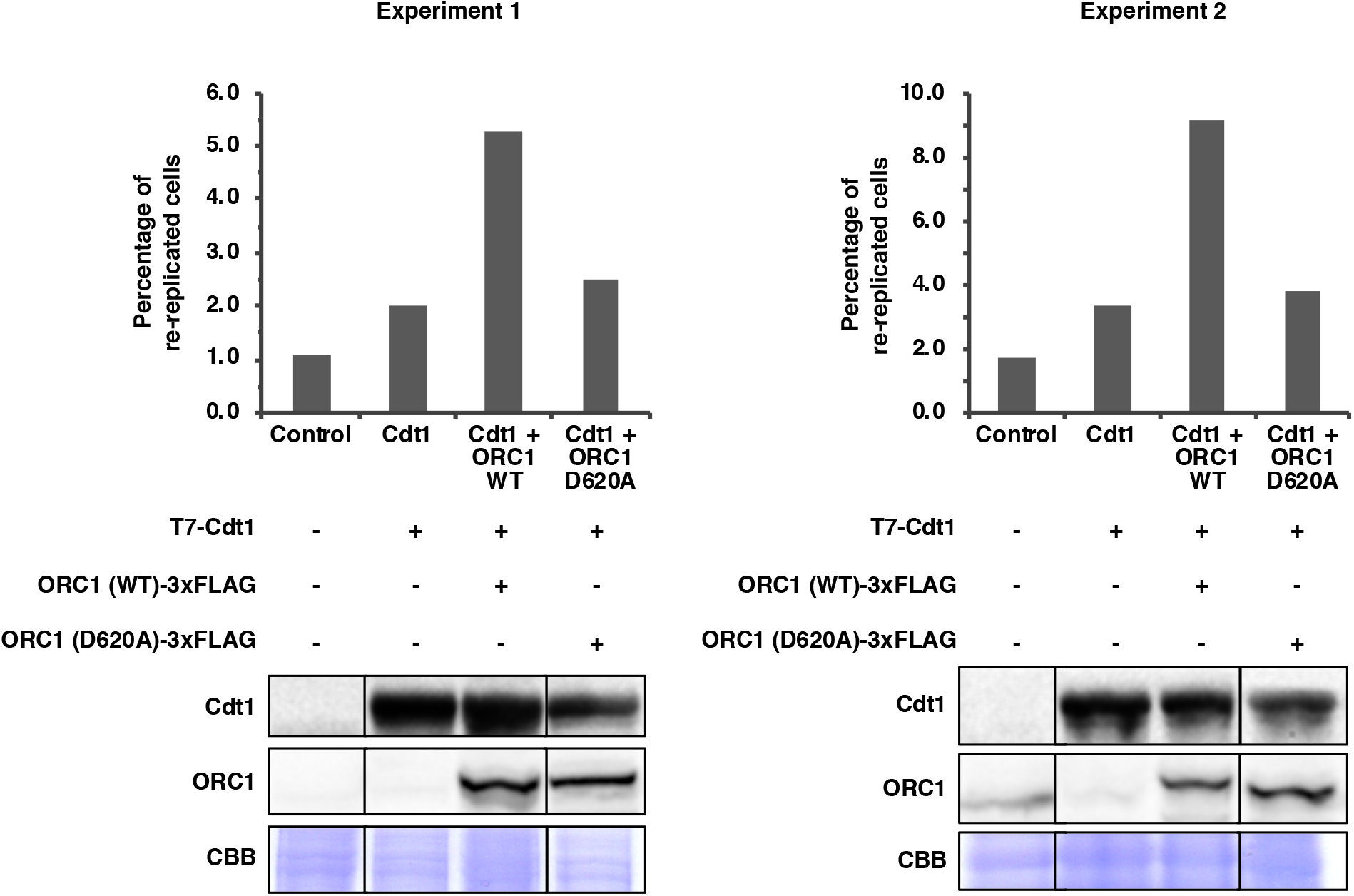
Re-replication induced by co-expression of Cdt1 + ORC1 is dependent on the ORC1 Walker B motif. HEK293T cells were co-transfected with a mixture of the expression vectors (T7-Cdt, ORC1 (WT)-3 × FLAG, and ORC1 (D620A)-3 × FLAG) or their empty vectors (-), as indicated for 48 h. *Top*, DNA content was analyzed by flow cytometry. The percentage of re-replicated cells was calculated as in Figure 5D. *Bottom,* whole-cell lysates were subjected to immunoblotting with the indicated antibodies. CBB staining serves as the loading control. The results of two independent experiments are shown. These data show that the disruption of ORC1 Walker B motif (D620A) abolishes re-replication induced by co-expression of Cdt1 + ORC1.

### Overexpression of ORC1 (244-511) compromises ORC and MCM recruitment to telomeres and allows telomeric DNA damage to accumulate under replication stress conditions

To investigate whether the overexpression of ORC1 (244-511) inhibits ORC binding to telomeres, HeLa cells stably overexpressing HA-ORC1 (244-511) were established by retroviral infection (Figure 6-figure supplement 1 A). Overexpression of HA-ORC1 (244-511) did not affect cell proliferation or cell cycle distribution (Figure 6-figure supplement 1 B and C). When ORC1-3×FLAG was transiently expressed in these cells to perform anti-FLAG ChIP analysis (Figure 6A), ORC1-3×FLAG binding at telomeres was significantly reduced upon overexpression of HA-ORC1 (244-511) (Figure 6B and Figure 6-figure supplement 2). Furthermore, overexpression of HA-ORC1 (244-511) significantly decreased MCM loading at telomeres (Figure 6C), while the total DNA co-precipitated with an anti-MCM7 antibody was unchanged (Figure 6D). These results suggest that HA-ORC1 (244-511) specifically impairs telomere binding of ORC and MCM. Using telomere-FISH and immunostaining of 53BP1, we next examined telomeric DNA damage upon overexpression of HA-ORC1 (244-511) (Figure 6E). In the absence of exogenous replication stress (Figure 6E; DDW), levels of spontaneous TIF-positive cells were low in both the control and HA-ORC1 (244-511)-overexpressing cells. When treated with 0.1 mM HU, cells overexpressing HA-ORC1 (244-511) showed a significant increase in TIF-positive cells compared with the control (Figure 6E; HU). These results further support the notion that TRF2-mediated ORC binding to telomeres contributes to telomere stability upon DNA replication stress.

**Figure 6.**
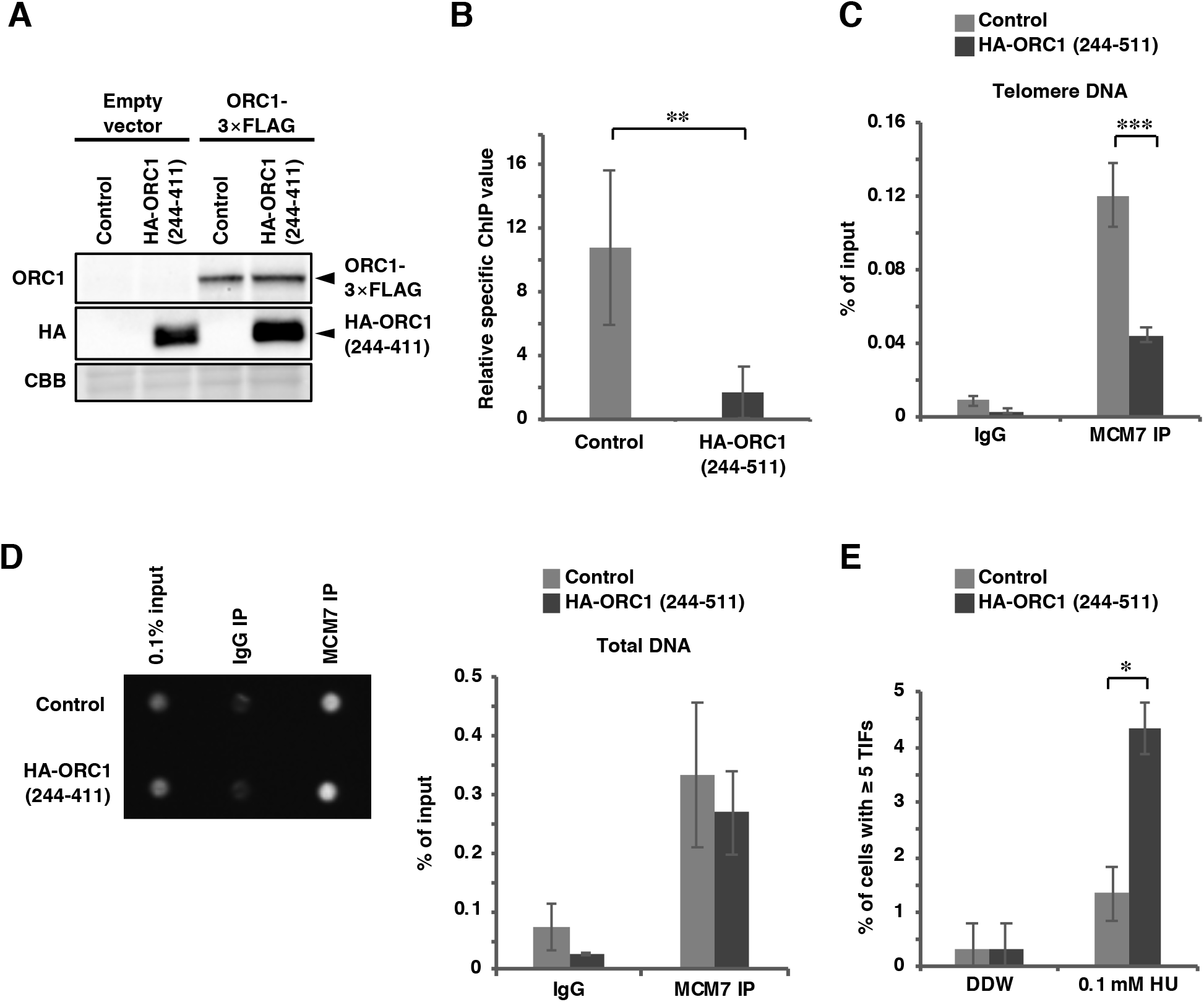
Overexpression of an ORC1 (244-511) fragment inhibits ORC- and MCM-binding to telomeres and exacerbates the accumulation of 53BP1 at telomeres upon replication stress. **(A and B)** Control and HA-ORC1 (244-511)-overexpressing HeLa cells were transfected with empty vector or ORC1-3 × Flag expression vector. At 48 h post-transfection, cells were subjected to ChIP with anti-FLAG antibody. (A) Cell lysates for ChIP assay were subjected to immunoblotting with the indicated antibodies. CBB serves as a loading control. (B) Purified DNAs from input and IPs were analyzed by qPCR with a primer pair amplifying a telomeric sequence or the *LMNB2* replication origin. Relative specific ChIP values were calculated as described in Figure 2E. The means ± SDs are shown (n = 6). **, P < 0.01 (two-tailed Student’s *t*-test). **(C and D)** Cells were subjected to the ChIP assay with control IgG or anti-MCM7 antibody. (C) Purified DNAs from input and IPs were analyzed by qPCR with a primer pair amplifying a telomeric sequence. Results are shown as the percent of input DNA. The means ± SDs are shown (n = 6). ***, P < 0.001 (two-tailed Student’s *t*-test). (D) *Left,* the total amounts of coprecipitated DNA and input DNA were analyzed using SYBR Gold staining and UV photography. *Right,* the means ± SDs are shown (n = 2). **(E)** Cells were treated with 0.1 mM hydroxyurea (HU) for 16 h. Telomere foci co-localizing with 53BP1 were analyzed as in Figure 3A and B. The frequency of cells carrying ≥5 TIFs was scored. The means ± SDs are shown (n = 3). One hundred cells were analyzed for each sample. The sum scores are used for statistical analyses. *, P < 0.05 (Fisher’s exact test).

**Figure 6–figure supplement 1.**
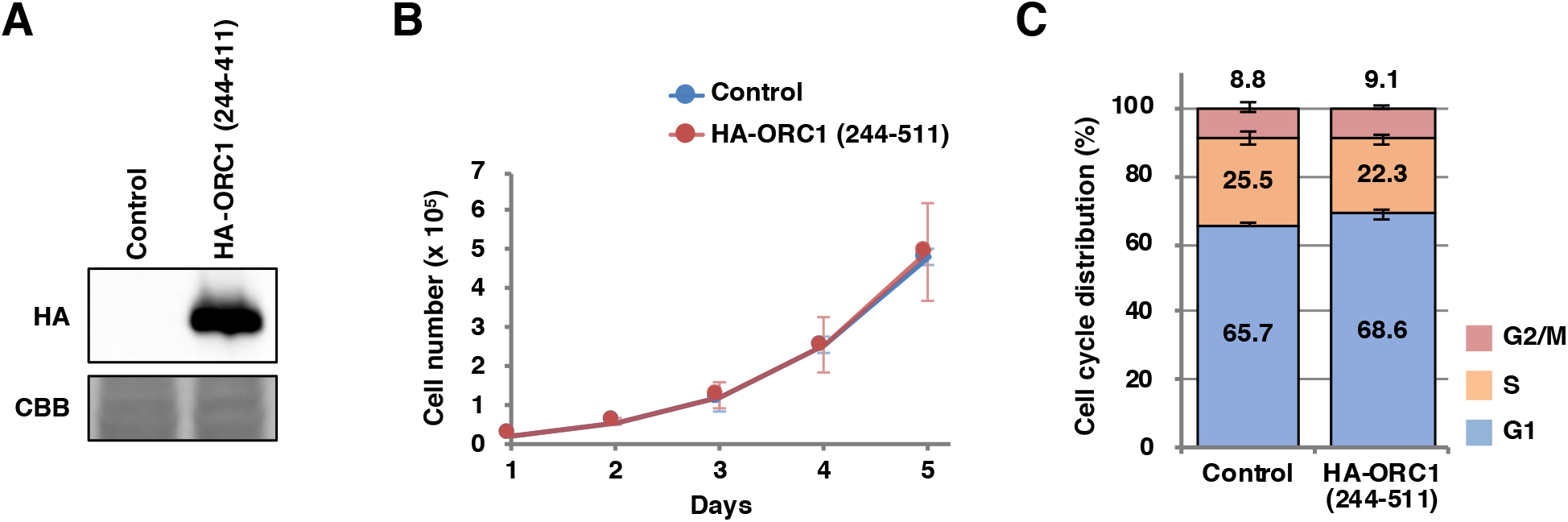
Overexpression of an HA-ORC1 (244-411) fragment does not affect HeLa cell proliferation. **(A)** HeLa cells stably expressing HA-ORC1 (244-411) fragment were established by retroviral infection. Expression of the introduced protein was analyzed by immunoblotting with anti-HA antibody. CBB serves as a loading control. **(B)** The cell growth was investigated for five days. The means ± SDs are shown (n = 2). **(C)** The cell cycle was analyzed by flow cytometry. The means ±SDs are shown (n = 2).

**Figure 6–figure supplement 2.**
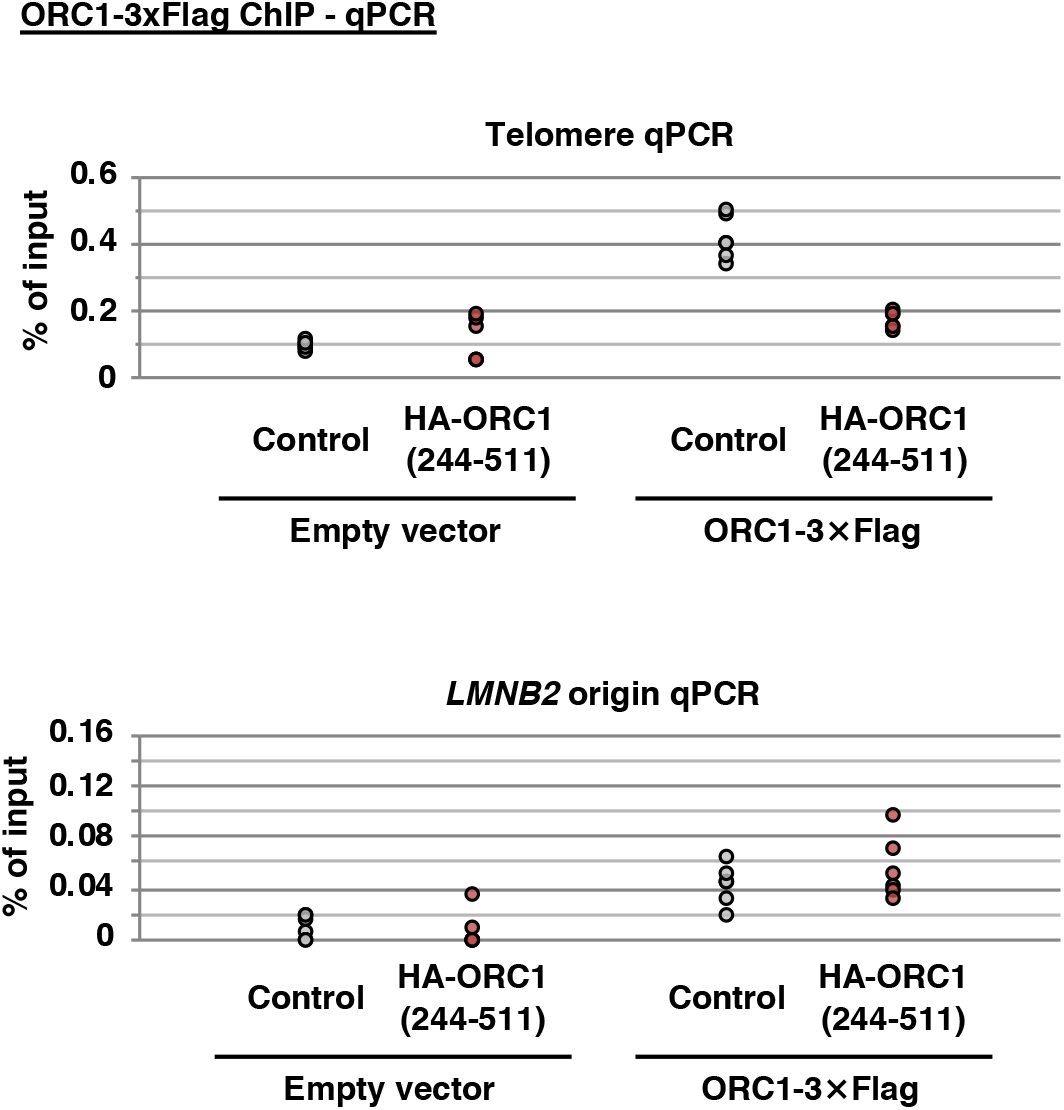
Source data of ChIP experiments. % of input values used for the calculation in Figure 6B are shown.

**Figure 6–figure supplement 3.**
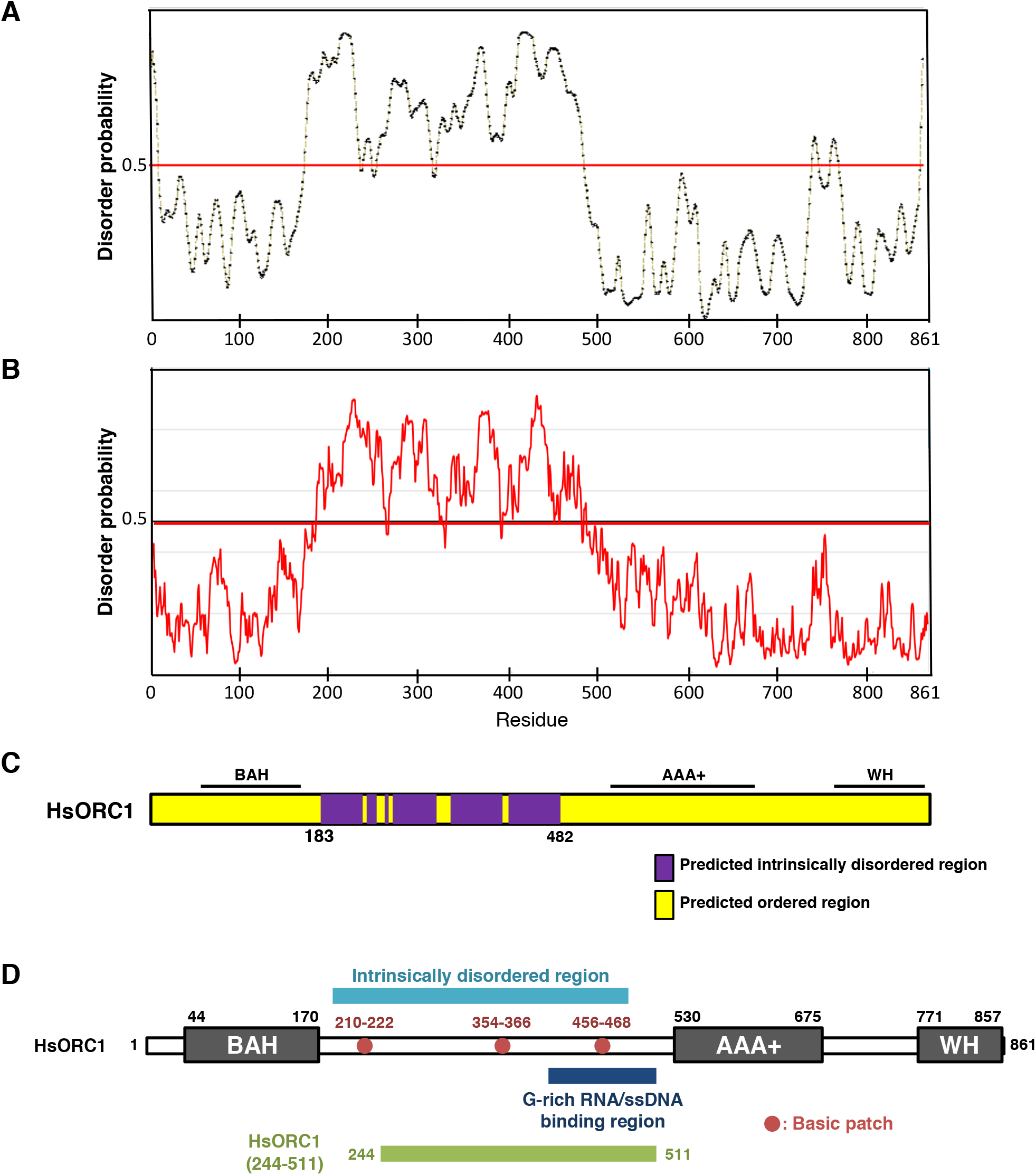
Schematics of ORC1 domain architecture showing the predicted intrinsically disordered region. **(A-C)** Intrinsically disordered region in ORC1 was predicted by (A) PrDOS (Ishida and Kinoshita, 2007) or (B) IUPred2A (Mészáros *et al.,* 2018). Predicted disorder probability is plotted. Red horizontal line represents the disorder/order threshold (a cutoff value 0.5) and residues scored above the line are predicted to be disordered. **(C)** Schematics of the intrinsically disordered region in ORC1. Purple rectangles represent overlapping regions of disorder predicted by each server. **(D)** Schematics of the full-length human ORC and ORC1 (244-511) fragments used in this study. BAH, bromo adjacent homology domain; AAA+, ATPases associated with diverse cellular activities domain; WH, winged-helix domain. References: 1. T. Ishida and K. Kinoshita (2007) PrDOS: Prediction of disordered protein regions from amino acid sequence. Nucleic Acids Res. 35: W460-W464. 2. B. Mészáros, G. Erdős, and Z. Dosztányi (2018) IUPred2A: Context-dependent prediction of protein disorder as a function of redox state and protein binding. Nucleic Acids Res. 46: W329–W337.

## Discussion

In this study, we specifically inhibited the TRF2-ORC interaction using the TRF2 E111A/E112A mutant, which is specifically defective for ORC1 binding (Figure 1). In HeLa TRF2 E111A/E112A knock-in clones, the recruitment of ORC and MCM to telomeres was reduced, and replication-stress-induced telomere instability was significantly enhanced (Figures 2 and 3). As an alternative experimental system, we identified an ORC1 fragment (244–511), which competitively inhibits the TRF2-ORC interaction. When this fragment was overexpressed in human cells, levels of telomere-bound ORC and MCM were reduced and replication stress-induced telomere DNA damage was enhanced (Figures 5 and 6). These specific inhibitions of the TRF2-ORC interaction avoid the limitations of simple TRF2 knockdown studies (Deng et al., 2007; Tatsumi et al., 2008), which result in the deprotection of telomere ends and prevent TRF2-binding factors from accessing telomeres. Our findings indicate a critical role for the TRF2-ORC interaction in the recruitment of ORC and MCM to telomeres and in the maintenance of telomere stability. It has recently been demonstrated that overexpression of a TRF2 ΔB mutant, an ORC-binding mutant that lacks the N-terminal basic domain (Atanasiu et al., 2006; Deng et al., 2007), decreases ORC recruitment to telomeres and leads to telomere fragility in LOX human melanoma cells (Drosopoulos et al., 2020). However, overexpression of TRF2 ΔB also induces cell cycle arrest, senescence, and rapid loss of telomeres (Deng et al., 2007; Drosopoulos et al., 2020; Saint-Léger et al., 2014; van Steensel et al., 1998; Wang et al., 2004). These phenotypes may be a consequence of the multiple roles the TRF2 basic domain has been shown to play in preventing homologous recombination-mediated telomere deletion (Poulet et al., 2009; Rai et al., 2016; Saint-Léger et al., 2014; Wang et al., 2004), promoting the replication of heterochromatin (Mendez-Bermudez et al., 2018), and mediating ORC recruitment at telomeres.

The dimerization of the TRF2 TRFH domain is required for the TRF2-ORC interaction (Higa et al., 2017b). Moreover, the TRF2 E111/E112 residues are critical for ORC recruitment (Figure 1). Our data suggest that the SLX4-binding region of TRF2 overlaps with the ORC1-binding region within the TRFH domain, yet the TRF2 E111A/E112A mutant retains its ability to interact with SLX4 while losing the ability to bind ORC (Figure 1B and 1F). In addition, the Y73, G74, V88, P90, K93, E94, H95, T96, S119, and M122 residues are also involved in ORC recruitment. The involvement of these residues, which are widely distributed, suggests that TRF2 and ORC1 interact through a broad binding interface. Indeed, we found that, although ORC1 (325–511) is sufficient for TRF2 binding, ORC1 (244–511) co-localizes with TRF2 with a 2-fold higher frequency (Figure 4). A flexible linker sequence is present between the N-terminal BAH (bromo adjacent homology) domain and the C-terminal AAA+ domain of ORC1 (Figure 6–figure supplement 3). This region is predicted to be a conserved, intrinsically disordered region (Figure 6–figure supplement 3) (Parker et al., 2019). Therefore, multiple residues in the ORC1 244–511 region may form a flexible binding interface for the interaction with TRF2.

Our results further reveal that TRF2 Y73A/G74A, V88A/P90A, E111A/E112A, and S119A/M122A mutations have no obvious effect on TRFH dimerization or the ability of TRF2 to bind Apollo, RAP1 or RETL1 (Figure 1C-E and G). TRF2 F120 is particularly important for the interaction with Apollo and SLX4 through their shared F/Y/HxLxP motif (Chen et al., 2008; Wan et al., 2013). RTEL1 binds to both TRF2 amino acids 64–83 and 312–341, with I79 playing a particularly crucial role in RTEL1 binding (Sarek et al., 2019, 2015). A RAP1-binding motif is found at the TRF2 275–316 region (Chen et al., 2011). The present data are in line with these findings. Although the TRF2 S119A/M122A mutations are in the proximity of F120, this mutant interacts with Apollo and SLX4, but not ORC1, suggesting that the TRF2 residues responsible for the bindings are different among these factors.

ChIP analysis revealed that ORC and MCM loading at telomeres is reduced by the overexpression of ORC1 (244–511) (Figure 6). Importantly, ORC1 (244–511) overexpression did not affect genome-wide DNA binding of MCM (Figure 6D), suggesting that ORC1 (244–511) overexpression selectively impairs the telomere binding activity of ORC. Moreover, re-replication induced by co-expression of Cdt1+ORC1 was not compromised by overexpression of ORC1 (244-511). The formation of the ORC complex should be unaffected by ORC1 (244-511) overexpression since the ORC1 N-terminal region (amino acids 1-470) is not essential for ORC complex formation (Tocilj et al., 2017). The BAH domain of ORC1 recognizes histone H4-K20me2 and regulates the loading of ORC onto replication origins, while mutations in this domain are implicated in the etiology of Meier-Gorlin syndrome (Beck et al., 2012; Bicknell et al., 2011; Kuo et al., 2012). The ORC1 244-511 region is situated away from the BAH domain and is unlikely to affect BAH domain-mediated binding to replication origins. The ORC1 244-511 region does, however, overlap with basic patches and G-rich RNA/ssDNA binding regions (Bleichert et al., 2018; Hoshina et al., 2013; Li et al., 2018) (Figure 6-figure supplement 3) thought to be involved in the interaction with origin DNA. It is possible that the binding of DNA through these regions of ORC1 is affected by ORC1 (244-511) overexpression. Future studies are required to evaluate the contribution of these regions to ORC recruitment to the G-rich telomere repeat sequence.

Dormant replication origins are critical for complete duplication of the genome (Alver et al., 2014; Moreno et al., 2016). A paucity of backup replication origins results in chromosome instability (Chuang et al., 2010; Ibarra et al., 2008; Kawabata et al., 2011; Shima et al., 2007). It has been reported that dormant telomeric origins are activated by replication stress to complete telomere replication (Drosopoulos et al., 2020). Our findings support the view that TRF2-mediated ORC and MCM recruitment plays a pivotal role in maintaining telomere stability through the formation of dormant telomeric origins that fire when replication forks stall or collapse.

## Materials and Methods

### Cell culture

U2OS, U2OS 2-6-3 (Higa et al., 2017b; Janicki et al., 2004), HEK293T, HeLa, TRF2-edited HeLa clones, and HCT116 cells were maintained in Dulbecco’s modified Eagle’s medium (Wako) supplemented with 8% fetal calf serum and antibiotics (0.1 mg/ml kanamycin).

### Plasmids

pSV40-HA-LacI, pSV40-TRF2-LacI, pSV40-TRF2 (45–244)-LacI, pSV40-TRF2ΔMyb-LacI, pcDNA3.1-zeo-ORC1-3×FLAG, pCLMSCV-HA-TRF2, and pCLMSCVhyg-T7-Cdt1 were described previously (Higa et al., 2017b; Sugimoto et al., 2008; Tatsumi et al., 2008, 2003).

pcDNA3.1-zeo-ORC1 (L229A)-3×FLAG, pcDNA3.1-zeo-ORC1 (D620A)-3×FLAG, pSV40-TRF2 (45–244/Y73A/G74A)-LacI, pSV40-TRF2 (45–244/V88A/P90A)-LacI, pSV40-TRF2 (45– 244/K93A/E94A/H95A/T96A)-LacI, pSV40-TRF2 (45–244/S98A/R102A)-LacI, pSV40-TRF2 (45– 244/E111A/E112A)-LacI, pSV40-TRF2 (45–244/S119A/M122A)-LacI, pENTR4-HA-TRF2 (Y73A/G74A), pENTR4-HA-TRF2 (V88A/P90A), pENTR4-HA-TRF2 (K93A/E94A/H95A/T96A), pENTR4-HA-TRF2 (S98A/R102A), pENTR4-HA-TRF2 (E111A/E112A), and pENTR4-HA-TRF2 (S119A/M122A) were produced by oligonucleotide-directed mutagenesis (Quick Change Site-directed Mutagenesis Kit; Stratagene) with the following mutagenic oligonucleotides and their complement.

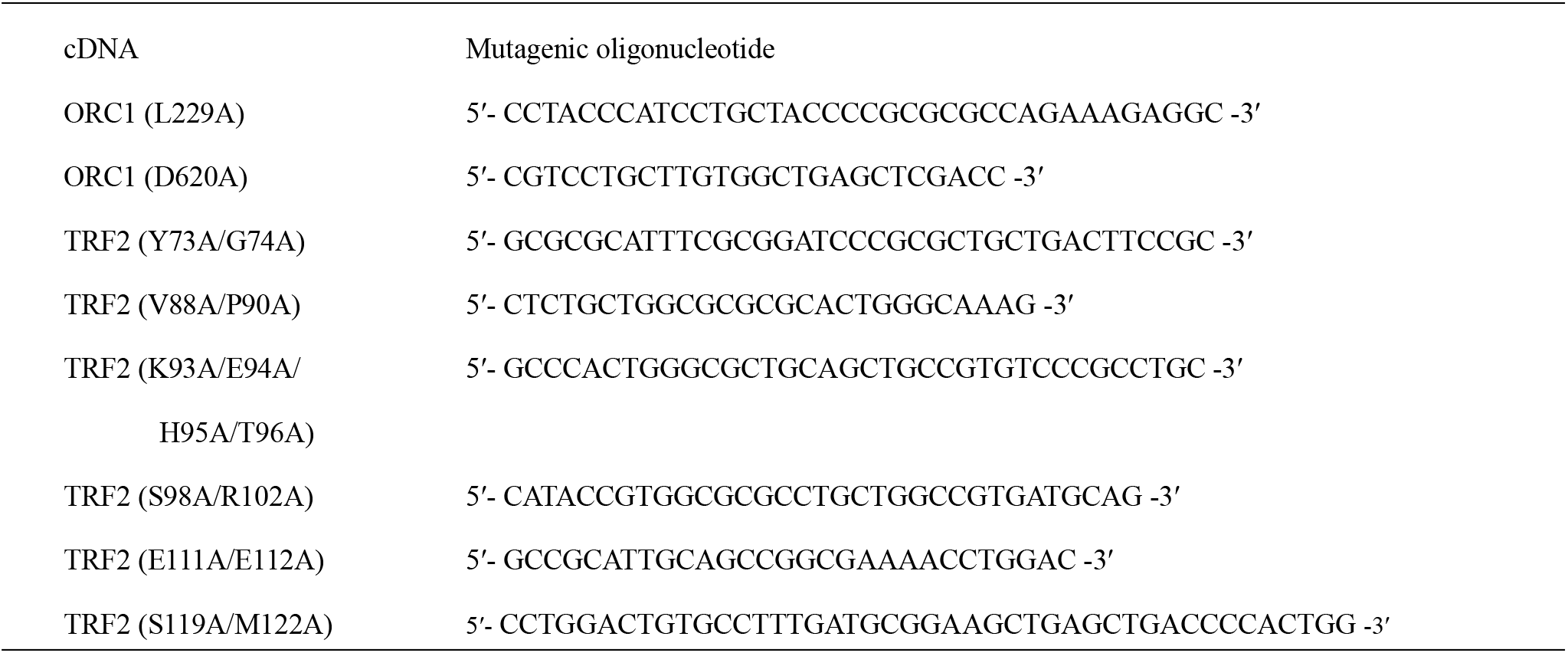

pAAVS1-CMV-HA-TRF2 and variants encoding alanine-substitution mutants of TRF2 (used in Figure 1E-G) were prepared using the In-Fusion (Clontech) reaction. The cDNAs were amplified by PCR with the primers listed below with a series of pENTR-HA-TRF2 vectors as template DNA. BglII- and SalI-digested pAAVS1-CMV (provided by Dr. Kanemaki, National Institute of Genetics, Mishima; Addgene # 105924) (Okumura et al., 2018) was used as the backbone vector.

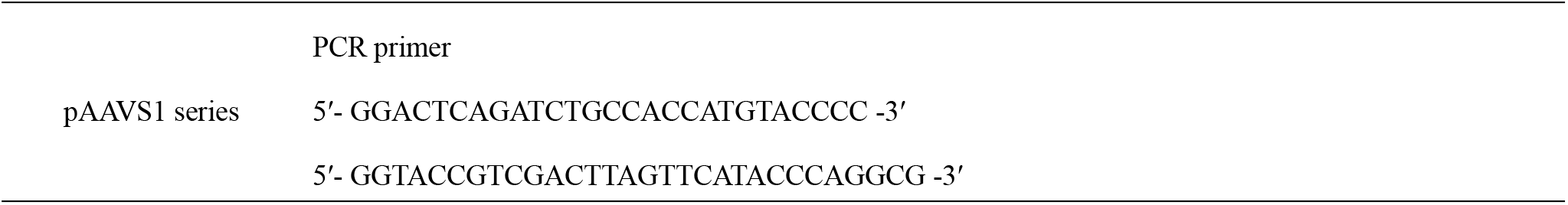

pFLAG-CMV-6a-Apollo was generated as follows: the cDNA was amplified by PCR from the template vector pCMV-SPORT6-Apollo (purchased from Dharmacon, clone# 5001181) using the primers listed below. Amplified cDNA and pFLAG-CMV-6a (SIGMA-ALDRICH) were digested by HindIII and PstI, and ligated using the Takara Ligation Kit (Takara, 6023).

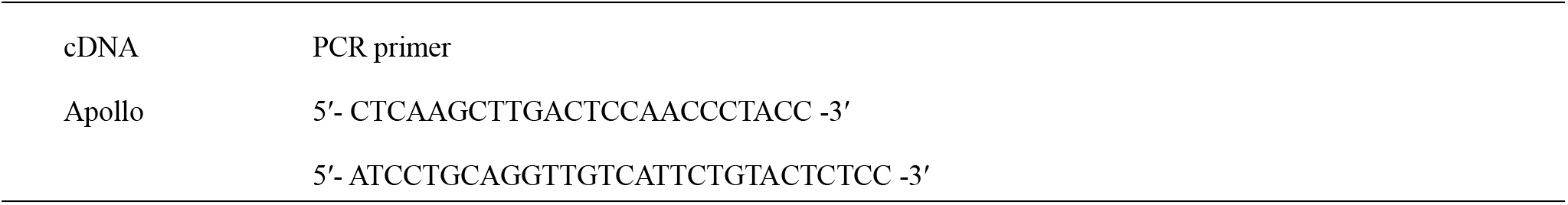

pFLAG-CMV-6a-SLX4 was generated as follows: the cDNA was amplified from pcDNA-FRT/TO GFP-BTBD12 (provided by Dr. Dario Alessi, University of Dundee, Scotland; clone number DU19216) (Muñoz et al., 2009) using the primers listed below. Amplified cDNA and HindIII-digested pFLAG-CMV-6a were subjected to the In-Fusion reaction.

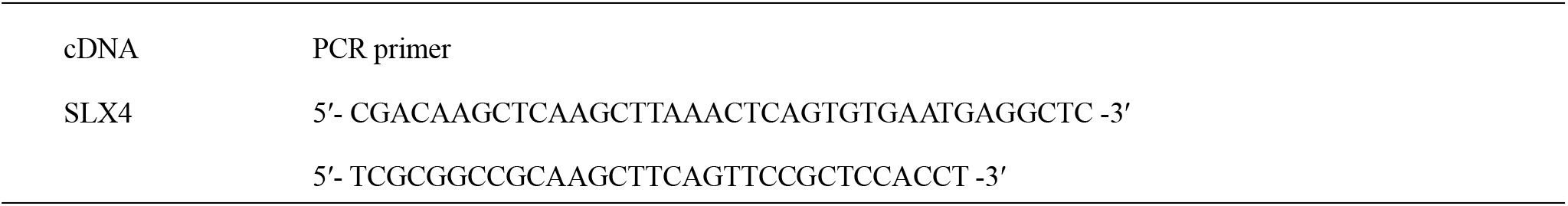

pFLAG-CMV-6a-ORC1 was prepared as follows: pBluescript-ORC1 (Fujita et al., 2002) was partially digested by NcoI, and a 5.5 kbp fragment was purified by agarose gel electrophoresis and gel extraction. After blunting using the Klenow fragment (Takara, 2140A), the fragment was further digested with NotI and ScaI, and a 2.4 kbp ORC1 fragment was purified by agarose gel electrophoresis and gel extraction. pFLAG-CMV-6a was digested with HindIII followed by blunting with the Klenow fragment, and further digested with NotI. After de-phosphorylation of the linear vector using Shrimp Alkaline Phosphatase (Takara, 2660A), the cDNA and the linear vector were mixed and ligated.

pX459-terf2-exon2-1 was generated by inserting a synthesized oligonucleotide into the pX459 vector (a gift from Feng Zhang, Broad Institute, Cambridge, MA; purchased from Addgene, # 62988) (Ran et al., 2013). To prepare the insert, the oligonucleotides listed below were mixed at final concentration of 4.5 μM each in 1× annealing buffer [20 mM Tris-HCl pH 7.5, 5 mM MgCl2, 30 mM NaCl, and 1 mM dithiothreitol (DTT)]. Mixed oligonucleotides were annealed by incubation at 100°C for 5 min followed by cooling at room temperature, and then phosphorylated by T4 polynucleotide kinase (Takara, 2021S). pX459 was digested with BbsI. Annealed oligonucleotides and linear vectors were mixed and ligated.

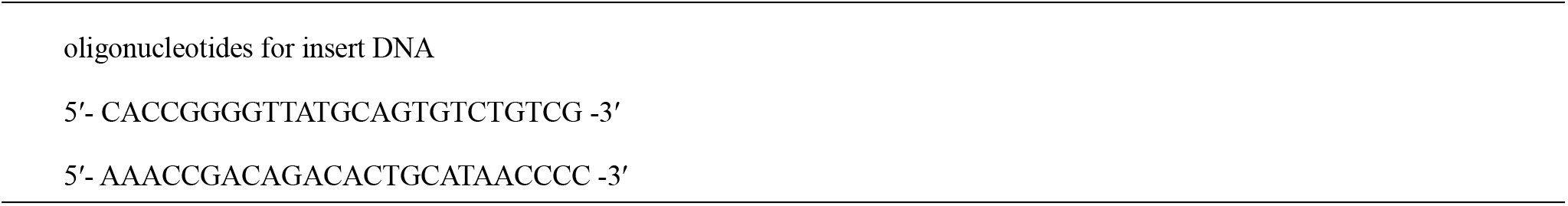

pFLAG-CMV-6a-ORC1 (2-511), pFLAG-CMV-6a-ORC1 (2-325), pFLAG-CMV-6a-ORC1 (2-244), and pFLAG-CMV-6a-ORC1 (2-85) were prepared by self-ligation of pFLAG-CMV-6a-ORC1 digested with the following restriction enzymes.

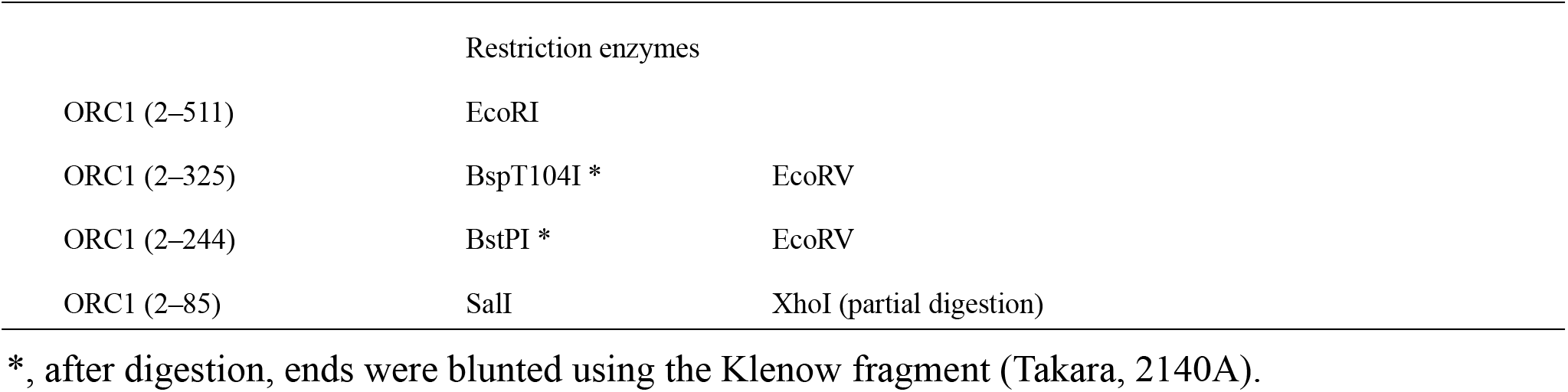

pFLAG-CMV-6b-ORC1 (244-511) and pFLAG-CMV-6b-ORC1 (325-511) were prepared as follows: pFLAG-CMV-6a-ORC1 was digested with BstPI [for ORC1 (244-511)] or BspT104I [for ORC1 (325-511)], blunted with the Klenow fragment, and further digested with EcoRI. pFLAG-CMV-6b (SIGMA-ALDRICH) was digested with HindIII, blunted with the Klenow fragment, and further digested with EcoRI. After de-phosphorylation of the linear vector using Shrimp Alkaline Phosphatase, the truncated ORC1 cDNA and the linear vector were mixed and ligated.

pFLAG-CMV-6a-ORC1 (Δ326-510), pFLAG-CMV-6a-ORC1(Δ386-510), pFLAG-CMV-6a-ORC1 (Δ411-510), pFLAG-CMV-6a-ORC1 (Δ446-510), and pFLAG-CMV-6a-ORC1 (Δ411-445) were prepared as follows: two fragments of ORC1 cDNA were amplified by PCR with the primers listed below using pcDNA3.1-zeo-ORC1-3×FLAG as a template. Amplified cDNA fragments and HindIII-digested pFLAG-CMV-6a were subjected to the In-Fusion or NEBuilder reaction (New England BioLabs, E2621).

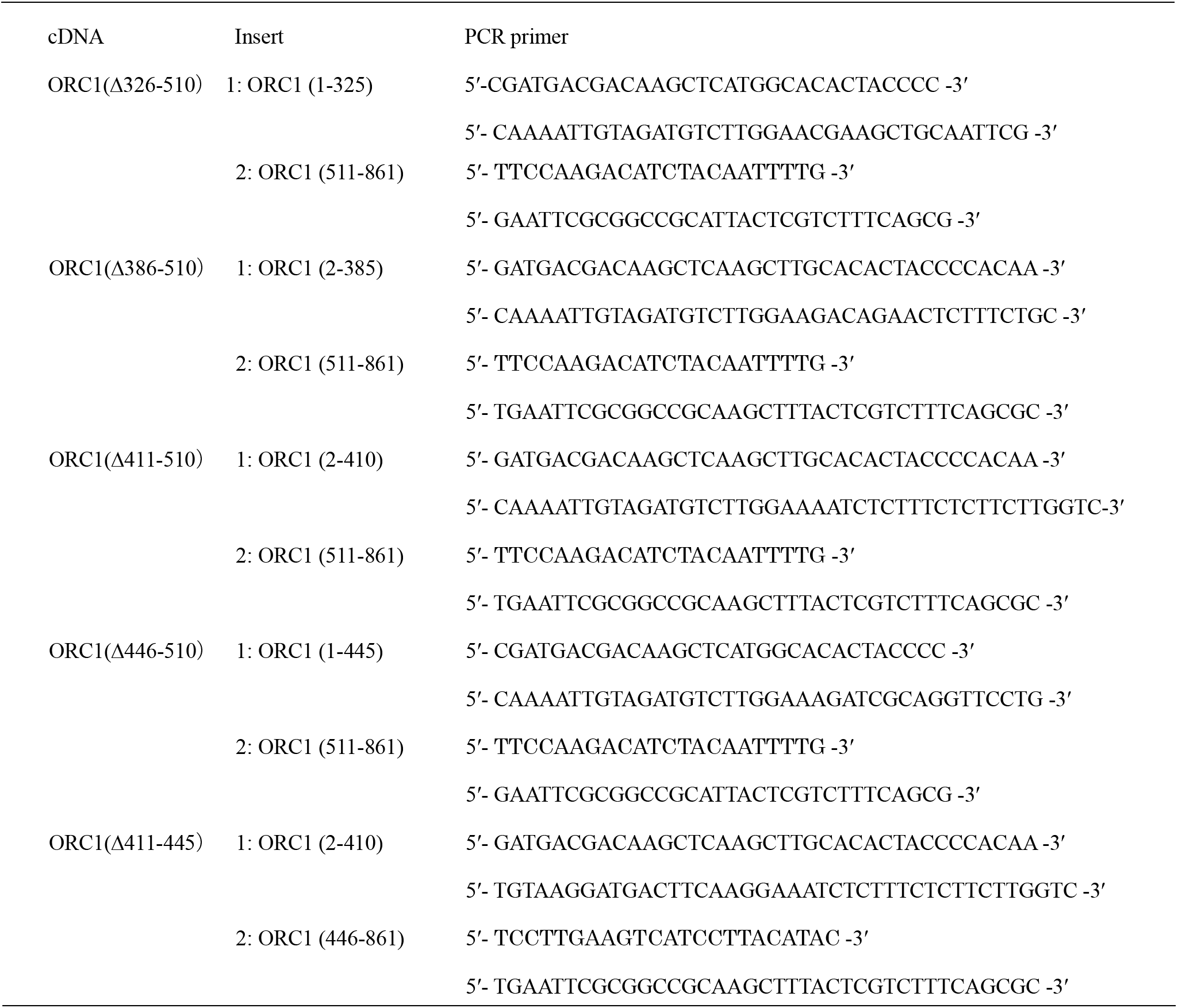

The retroviral expression vector pQCXIP-HA-ORC1(244-511) was prepared by In-Fusion reaction. The cDNAs were amplified by PCR with the primers for N-terminal HA-tagging listed below using pcDNA3.1-zeo-ORC1-3×FLAG as template DNA. NotI-digested pQCXIP (Clontech) was used as the backbone vector.

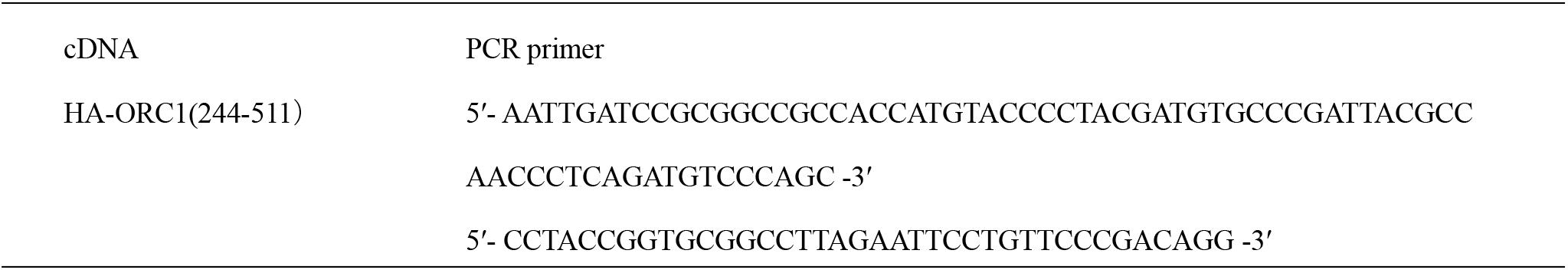

As described previously (Higa et al., 2017b; Tatsumi et al., 2008), our TRF2 cDNA was chemically synthesized with codon optimization and thus some oligonucleotide sequences described here are not comparable to those of cell-derived TRF2 cDNA. The length of our TRF2 oligopeptide is 500 amino acids, which is 42 amino acids shorter at the N-terminus (542 amino acids in total) than that used in studies of the RTEL1-binding regions of TRF2 (Sarek et al., 2019, 2015). Therefore, the apparent position of residues differ between these studies.

### Transfection

For immunofluorescence analysis, expression plasmids (total: 0.56 μg for Figs. 1B, 1D, 4B, 4D; 0.84 μg for Fig. 5B) were transiently transfected into 8 × 10^4^ U2OS 2-6-3 cells in 4-well chamber slides using Lipofectamine 2000 reagent (Invitrogen, 11668019) according to the manufacturer’s instructions.

For immunoprecipitation, ChIP and re-replication analysis, transfection was performed with PEImax reagent (Polysciences, 24765), as previously described (Sugimoto et al., 2015). Detailed transfection conditions used in the experiments are listed below. For ChIP analysis (Figures 2E, F, and 6A, B), the medium was changed 6 h after transfection.

| Experiment | Cells | Total DNA | Cellular density |
| --- | --- | --- | --- |
| Fig. 1C | U2OS | 2 μg | $5 \times 10^5$/6 cm dish |
| Fig. 1E | HCT116 | 4 μg | $1 \times 10^6$/6 cm dish |
| Fig. 1F | HEK293T | 3 μg | $3 \times 10^6$/6 cm dish |
| Fig. 1G | HEK293T | 8 μg | $6 \times 10^6$/10 cm dish |
| Fig. 2E and F | TRF2-edited HeLa clones | 19 μg | $3.8 \times 10^6$/15 cm dish |
| Fig. 5A | HEK293T | 18 μg | $6 \times 10^6$/10 cm dish |
| Fig. 5C-E | HEK293T | 4 μg | $2 \times 10^6$/6 cm dish |
| Fig. 6A and B | HeLa expressing ORC (244-511) | 7.5 μg | $1.5 \times 10^6$/10 cm dish |

### Immunofluorescence staining

For Figure 1B, co-staining of endogenous ORC1 with LacI were performed as described previously (Higa et al., 2017b). Briefly, cells were washed with PBS (Wako, 048-29805), fixed with chilled 100% methanol for 10 min, permeabilized with 0.1% Triton X-100 in PBS for 10 min, and then immunostained.

For the staining of FLAG-Apollo, and FLAG-ORC1 truncation mutants (Figures 1D and 4B), cells were fixed with 3.7% formaldehyde (Nacalai Tesque, 16223-55) in PBS. For the staining of full-length FLAG-ORC1 and FLAG-ORC deletion mutants (Figures 4D and 5B), cells were fixed with chilled 100% methanol for 10 min, and permeabilized with 0.1% Triton X-100 in PBS for 10 min. The staining of 53BP1 was performed after PNA-FISH (Figures 3A, B and 6E). Cells were then incubated with primary antibodies in PBS supplemented with 1% fetal calf serum for 1 h at room temperature for 53BP1 staining or 2 h at 37°C for LacI co-staining, followed by incubation with secondary antibody for 1 h at room temperature and 0.2 μg/ml 4,6-diamidino-2-phenylindole (DAPI) in PBS for 15 min at room temperature. Cells were washed three times after each staining with PBS for 5 min at room temperature, and finally mounted in Fluoro-KEEPER Antifade Reagent (Nacalai Tesque, 12593-64) and stored at 4°C.

Microscopic analysis was performed with the KEYENCE BZ-9000 or BZ-X700 fluorescence microscope. Co-localization frequency of foci of each protein with the LacI foci was examined. A summary of the co-localization frequency calculated from multiple biologically independent experiments is shown in the figures. The values are sum scores from at least two independent experiments performed for each condition. At least 31 cells with a prominent LacI focus were scored for each experiment/condition.

### Immunoprecipitation

For Figure 1C, E and F, cells were lysed in 500 mM NaCl NET gel buffer (500 mM NaCl, 0.1% Triton X-100, 50 mM Tris-HCl pH 7.4). The lysates were diluted with 0 M NaCl NET gel buffer to a final NaCl concentration of 150 mM and then subjected to immunoprecipitation with 2 μg antibody. Bound proteins were eluted in 1× sample buffer (62.5 mM Tris-HCl pH 6.8, 2% SDS, 5% 2-mercaptoethanol, 10% glycerol, 0.01% bromo-phenol blue).

For Figures 1G and 5A, transfected HEK293T cells were cross-linked with 1% formaldehyde and then subjected to immunoprecipitation as described previously (Higa et al., 2017b) using 2 μg (Fig. 1G) or 4 μg (Fig. 5A) of antibody.

### Immunoblotting and antibodies

Immunoblotting was performed as described previously (Sugimoto et al., 2008). Coomassie brilliant blue (CBB) staining was performed with the Rapid Stain CBB Kit (Nacalai Tesque, 30035-14). Quantification was performed with the LumiVision Analyzer (for immunoblots, AISIN Seiki) or ImageJ software (for CBB, NIH).

Preparation of polyclonal rabbit antibodies against human ORC1, Cdt1 and MCM7 was described previously (Tatsumi et al., 2008). Rabbit anti-LacI antibody was obtained by immunizing rabbits with a bacterially produced His-T7-LacI protein.

Other antibodies used in this study are as follows: LacI (clone 9A5, Merck, 05-503), ORC1 (Santa Cruz, ac-23887), HA-tag (clone 3F10, Roche), HA-tag (COVANCE MMS-101R), FLAG-tag (clone M2, SIGMA-ALDRICH), FLAG-tag (Thermo, PA1-984B), RAP1 (Bethyl, A300-306A), RTEL1 (Novus, NBP2-22360), SLX4 (Novus, NBP1-28680), TRF2 (Merck, 4A794), 53BP1 (Novus, NB100-904), rabbit normal IgG (DAKO, X0903), mouse normal IgG (Southern Biotech, 0107-01), horseradish-peroxidase (HRP)-conjugated anti-mouse IgG (H + L) (Invitrogen, 61-6520 or ROCKLAND, 18-8817-33), HRP-conjugated anti-rabbit IgG (H + L) (Invitrogen, 65-6120 or ROCKLAND,18-8816-33), HRP-conjugated anti-rat IgG (H + L) (Zymed, 62-9520), CF488A-conjugated goat anti-rabbit IgG (Biotium, 20019), CF594-conjugated donkey anti-rabbit IgG (Biotium 20152), CF594-conjugated goat anti-mouse IgG (Biotium, 20111), Alexa488-conjugated goat antimouse IgG (Invitrogen, A11029).

### Cas9-based gene-editing

Cas9-based gene-editing was performed by transient transduction of Cas9/guide RNA and ssODN (Ran et al., 2013). HeLa cells cultured in 6 well plates (1×10^5^/well) were co-transfected with 2 μg pX459-terf2-exon2-1 and 2 μg ssODN using Lipofectamine 2000 reagent. ssODN carries the following sequence: ssODN WT, 5’-AGAGAAGAACACAAAAATAGCCATACCTAAATTTTCCCCTTCTTCAATTCGCGACAGACACTGCATAACCCGCAGCAATCGGGACACGGT-3’; ssODN EE, 5’-AGAGAAGAACACAAAAATAGCCATACCTAAATTTTCCCCTGCTGCAATTCGCGACAGACACTGCATAACCCGCAGCAATCGGGACACGGT-3’.

After puromycin (0.5 μg/ml) selection for 3 days, cells were subjected to limiting dilution. Aliquots of the obtained clones were re-seeded into 96-well plates, and clones were screened by RFLP analysis. Positive clones were re-subjected to further limiting dilution. After the second selection, confirmed positive clones by RFLP analysis were used for subsequent experiments.

### RFLP analysis

For screening of HeLa clones, cells were seeded into 96-well plate and rinsed with PBS once after reaching confluence. Forty-five microliters of 50 mM NaOH were then added to each well, and the cells were lysed in a 90–95°C water bath. Cell lysates were directly added to the PCR reaction mix (Toyobo, KOD-201).

For Figure 2B, genomic DNA was prepared as follows: cells were lysed with SDS and treated with RNase A (Invitrogen) and proteinase K (Roche, 3115887001). Genomic DNA was precipitated by isopropanol (Nacalai tesque, 29113-95). Purified genomic DNA was dissolved in TE (10 mM Tris-HCl pH 8.0 and 1 mM EDTA) and fragmented by incubation at 70°C for 2–4 h.

The following primers were used to amplify the target site in *TERF2* gene exon 2: 5’-GGACTTCAGACAGATCCGGG-3’ and 5’-CTCCTCAGATACGAGTGGCAAG-3’. Amplified DNA was incubated with or without NruI and separated by agarose gel electrophoresis.

### Sequencing of the target locus of the *TERF2* gene

The target sites in the *TERF2* gene exon2 were amplified as described above. For TA cloning, the PCR products were 1000-fold diluted and re-amplified with Taq polymerase (Bioacademia, 02-001) and the same primer pair. The products were purified by agarose gel extraction, cloned into T-Vector pMD20 (Takara, 3270), and sequenced (Thermo, 4337454). Eleven colonies from each clone were analyzed.

### Chromatin immunoprecipitation and quantitative real-time PCR (qPCR) analysis

Chromatin immunoprecipitation was performed as described previously (Ishimoto et al., 2021). For qPCR analysis, SYBR Premix Ex Taq II (Takara, RR081A) or TB Green Premix Ex Taq II (Takara, RR820A) was used according to the manufacturer’s instructions. PCR reactions were performed using the CFX96 Touch Real-Time PCR Detection System (BIO-RAD). For detection of the telomere sequences and *LMNB2* replication origin region, the primer sequences and qPCR cycling parameters were as described previously (Cawthon, 2002; Higa et al., 2017b; Sugimoto et al., 2011). The total amount of co-precipitated DNA was quantified using SYBR Gold staining (Invitrogen, S11494) (Sugimoto et al., 2015). SYBR Gold signals were captured using cooled-CCD camera detection systems (LumiVision Imager, AISIN Seiki) within the linear range, and band intensities were quantitated using LumiVision Analyzer.

### Fluorescence-activated cell sorting (FACS)

Cells were trypsinized and suspended in PBS supplemented with 0.1% Triton X-100 and 10 μg/ml RNase A. After staining with propidium iodide (40 μg/ml), cell cycle distribution was examined using a FACS Verse flow cytometer (BD Bioscience) and ModiFit LT (Verity Software House). For Figure 5 D and E, re-replication was measured as described previously (Sugimoto et al., 2009). Dots with higher FL2-W signals, which result from aggregated cells and cell debris, were excluded from measurements of re-replication.

### PNA-FISH: fluorescence in situ hybridization with peptide nucleic acid probes

After rinsing with PBS, cells were fixed with 3.7% formaldehyde (for Figures 3A, B and 6E) or chilled 100% MeOH (for Figure 3C and D). For formaldehyde fixation, cells were permeabilized in PBS containing 0.1% TX-100. Cells were dehydrated by incubating in serially diluted EtOH (70%, 95%, and 100%). After drying cells completely, hybridizing solution [70% formamide, 9 mM Tris-HCl pH 7.5, 1% skimmed milk, and 200 nM Cy5-TelC (Panagene, F1003)] was added to each well. After placing coverslips onto slides, they were baked at 80°C for 5 min, followed by incubation at room temperature overnight. After removing coverslips, cells were washed once with PBS, twice with washing solution (70% formamide and 9 mM Tris-HCl pH 7.5) at room temperature for 15 min, and three times with PBS. For co-staining with 53BP1, immunofluorescence staining was performed after PNA-FISH.

### Establishment of HeLa cells stably expressing HA-ORC1 (244-511)

HeLa cells were infected with the recombinant retroviruses encoding HA-ORC1(244-511) or an empty vector, as described previously (Sugimoto et al., 2009). Infected cells were selected with puromycin (0.5 μg/ml) and subjected to assays without cloning.

### Data presentation and statistical analysis

Unless otherwise stated, quantitative data are represented as the mean ± SD of three or more independent experiments. The number of experiments was chosen according to the standards of the field. The statistical analyses used are defined in each figure legend. For Figures 1B, 1D, 3B, 4B, 4D, 5B, and 6E, the sum scores of two or more independent experiments were used for the statistical analyses. Individual values of each experiment are provided as Source data files. Only for Figure 4–figure supplement 1, the values represent the score from a single experiment. For the *t*-tests, data distribution was assumed to be normal but this was not formally tested. Tukey-Kramer multiple comparison test was performed after examination by one-way ANOVA. P-values less than 0.05 were considered statistically significant. For qualitative data and semi-quantitative data, a representative image from multiple independent experiments is shown; for all such figures, essentially the same results were obtained in the multiple independent experiments.

## List of Figures, figure supplements, and source data

Figure 1, Figure 1-source data 1

Figure 2, Figure 2–figure supplement 1, Figure 2-source data 1

Figure 3, Figure 3-source data 1

Figure 4, Figure 4–figure supplement 1, Figure 4-source data 1 Figure 5, Figure 5–figure supplement 1, Figure 5-source data 1

Figure 6, Figure 6–figure supplement 1, Figure 6–figure supplement 2, Figure 6–figure supplement 3, Figure 6-source data 1

## Acknowledgements

We thank Tohru Kiyono (National Cancer Center EPOC, Kashiwa), Masato Kanemaki (National Institute of Genetics, Mishima), Yoko Katsuki (Kyoto University, Kyoto), Minoru Takata (Kyoto University, Kyoto), Hilary McLauchlan (University of Dundee, Scotland), and John Rouse (University of Dundee, Scotland) for plasmids. We thank Toshiki Tsurimoto (Kyushu University, Fukuoka), Tsutomu Katayama (Kyushu University, Fukuoka), and Hironori Kawakami (Sanyo-Onoda City University, Sanyo-Onoda) for discussion and comment. We appreciate technical support from the Research Support Center, Research Center for Human Disease Modeling, Kyushu University Graduate School of Medical Sciences. This investigation was supported, in part, by a grant to KY from the Japan Society for the Promotion of Science (KAKENHI grant number JP17K15065). MH was supported by a JSPS research fellowship (JP18J11443).

## Author contributions

KY and MF designed and managed the project. MH, YM, JY and KY conducted the experiments and data analysis. NS and MF supervised the experiments and data analysis. MH, KY, and MF wrote the manuscript.

